# Multimodal Neuroimaging Reveals Distinct Characteristics of Levodopa-Induced Dyskinesias in *de novo* Parkinson’s Disease Patients

**DOI:** 10.1101/2025.01.20.633873

**Authors:** Sakshi Shukla, Mantosh Patnaik, Aditya Kumar, Sule Tinaz, Nivethida Thirugnanasambandam

**Author notes:** These authors jointly supervised this work (Sule Tinaz, Nivethida Thirugnanasambandam). Corresponding author: Nivethida Thirugnanasambandam.

## Abstract

Levodopa-induced dyskinesia (LID) is a significant treatment complication that affects a substantial proportion of Parkinson’s disease (PD) patients. Our understanding of the neural basis of LID remains limited, partly due to the small sample sizes in existing neuroimaging studies.

In this study, we utilized structural MRI data from the Parkinson’s Progression Markers Initiative (PPMI) database, including de novo PD patients (104 non-dyskinetic for a least 3 years after diagnosis and 120 who developed dyskinesia) and 100 age- and sex-matched healthy controls. Additionally, we analyzed resting-state functional MRI data from a subset of these participants to investigate connectivity differences among the groups.

Our analysis revealed no significant baseline volumetric differences between dyskinetic and non-dyskinetic PD patients. However, the thickness of frontal and sensorimotor cortices were significantly greater in dyskinetic patients. In the subcortical regions, vertex-based shape analysis identified localized surface growth in the left caudate and left pallidum, as well as surface morphology changes in the bilateral pallidum in dyskinetics. Resting-state functional connectivity analysis revealed stronger connectivity between the putamen, inferior frontal gyrus, and sensory cortex in dyskinetic PD patients compared to non-dyskinetics.

These findings suggest that specific morphological and functional changes in the motor cortical-basal ganglia circuitry of de novo PD patients may predispose them to LID over time. Additionally, the altered functional connectivity patterns reinstate the role of the inferior frontal gyrus in the pathophysiology of dyskinesia and suggest that it might be a suitable target for neuromodulatory interventions, consistent with previous reports.

## Introduction

Idiopathic Parkinson’s disease (PD) is a multifaceted neurodegenerative disorder manifesting with both motor and non-motor symptoms. Patients are usually diagnosed when nearly 50-60% of dopaminergic neurons in the substantia nigra have degenerated (J M Fearnley and A J Lees 1991). Dopamine replacement therapy with levodopa is currently the most effective strategy to alleviate motor symptoms. However, in about 40% of patients, chronic levodopa intake results in motor side effects within 4-6 years (Ahlskog et al., 2001), known as levodopa-induced dyskinesias (LID) (Yahr et al., 1968). Dyskinetic patients exhibit involuntary dystonic and/or choreic movements, which can be more debilitating than freezing of gait (Thanvi B et al., 2007). Young-onset disease, more severe disease, and higher doses of levodopa have been identified as risk factors for LID (Kumar N et al., 2005, Di Monte DA et al., 2000, Nutt JG et al., 1992). The benefits and chronic side effects of levodopa can be balanced to some extent by carefully tailoring its dosage. The most effective treatment currently available for LID is deep brain stimulation, which allows for the reduction of levodopa dosage. However, it is a surgical option and may not be suitable for all patients (Martini et al., 2019). It would, therefore, be beneficial to identify patients with a higher risk of developing LID earlier in the course of the disease, so that treatment strategies can be personalized to minimize or delay potential disability.

Neuroimaging studies have contributed to our understanding of LID pathophysiology indicating the involvement of basal ganglia-cortical circuits (Cerasa et al., 2012, Beaudoin-Gobert et al., 2018). For example, a significant increase in the grey matter volume of bilateral inferior frontal gyri in dyskinetic compared to non-dyskinetic PD patients has been reported (Cerasa et al., 2011). This increase was negatively correlated with age at disease onset, suggesting that remodeling of the prefrontal cortex over time in younger patients might play a role in the pathophysiology of LID. In a later study, an increase specifically in the cortical thickness of the right inferior frontal sulcus in dyskinetic PD patients was shown (Cerasa et al., 2013). On the other hand, a recent MRI study in a rat model of LID revealed bilateral volume reduction in subcortical and cortical areas (Zhang et al., 2021).

Studies using mere volumetric analyses have reported conflicting results, some showing reduced striatal volume (Geng et al., 2006, Garg et al., 2015, Oltra et al., 2022), and others showing no difference in striatal volume (Nemmi et al., 2015, Khan et al., 2019, Gong et al., 2020) in PD patients compared to healthy controls. The picture is even less clear when comparing the volumetric changes in dyskinetic versus non-dyskinetic PD patients. However, surface-based shape analysis can detect subtle subcortical structural abnormality. For example, in a recent study (Youn et al., 2022), no significant volumetric differences were found in the basal ganglia structures in the dyskinetic compared to the non-dyskinetic PD group. Yet, shape analysis revealed local atrophy in the right globus pallidus internus only in the dyskinetic PD group. This suggests that surface-based shape analysis can provide a deeper insight into the regional specificity of volumetric changes. Notably, all structural MRI investigations concerning dyskinesias have focused solely on patients *after* the development of LID. The cortical and subcortical volumetric and shape alterations in a cohort of de novo, unmedicated PD patients who later developed LID have not been explored thus far. Examining the brains of this cohort will shed light on whether specific structural changes in the brain render some patients more prone or resistant to LID.

Functional MRI (fMRI) studies have demonstrated that levodopa treatment modulates the resting-state functional connectivity (FC) of the sensorimotor network in PD patients (Cerasa et al., 2015, Esposito et al., 2013). A landmark study showed an accurate prediction of dyskinesia in PD patients based on the resting-state FC between putamen and sensorimotor cortex (Herz et al., 2016). However, the relatively small sample size (N=10-15) is a limitation in these studies. It is only recently that patient databases such as the Parkinson’s Progression Markers Initiative (PPMI) have enabled the analysis of large datasets to arrive at more robust results (Marek et al., 2018).

In the current study, we curated structural MRI and fMRI scans from the PPMI repository. We aimed to examine MRI data from a relatively large number of *de novo* PD patients and identify distinct MRI features that can differentiate those who subsequently developed LID from those who did not. We used volumetric, surface-based shape analysis along with resting-state FC methods for this purpose. We hypothesized that PD patients who eventually developed LID would demonstrate different morphometry and patterns of FC in the frontal/sensorimotor cortex-basal ganglia network even at baseline or before LID onset compared to those who did not develop LID.

## Materials and Methods

### Participants

#### Structural MRI data

We downloaded the T1-weighted (T1w) MRI scans of 343 subjects and their metadata from the PPMI website (https://www.ppmi-info.org/) on January 27, 2022 (Parkinson Progression Marker Initiative 2011). After quality checks, 19 subjects were excluded from the original cohort due to significant artifacts in the original T1w image and improper segmentation during preprocessing (see **Figure 1** for the overall pipeline).

**Figure 1:**
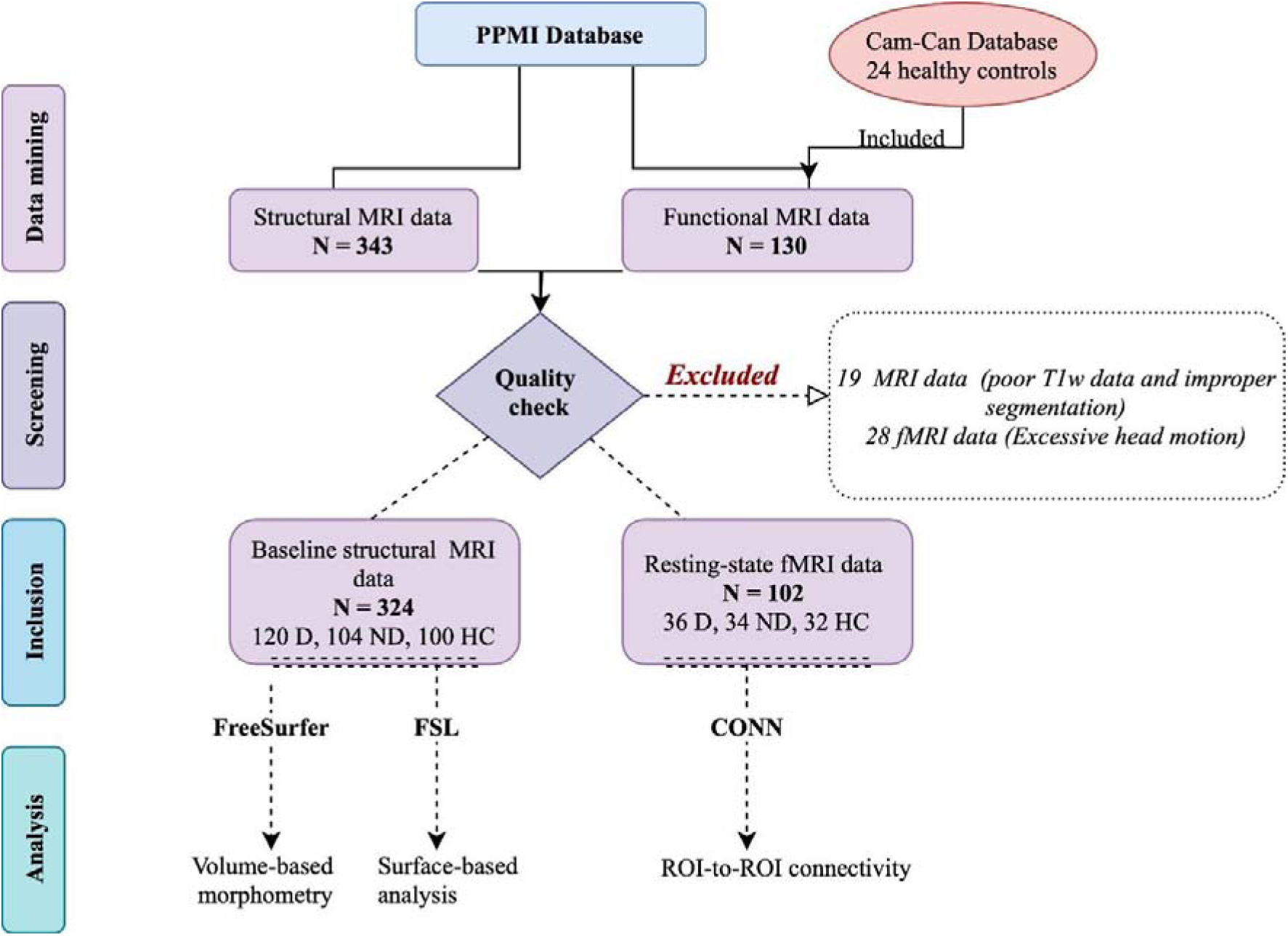
Study pipeline. D: Dyskinetics, ND: Non-dyskinetics, HC: Healthy Controls, PPMI: Parkinson’s Progression Markers Initiative, FSL: FMRIB software library, CONN: Connectivity toolbox.

The protocols for PPMI data acquisition have been provided in the PPMI’s imaging technical operation manuals (www.ppmiinfo.org). All T1w structural and T2*w functional MRI scans included in this study were acquired in 3T scanners (see supplementary material Table S1 & S2 for detailed MRI parameters of the PPMI and Cam-CAN datasets (Taylor J.R. et al., 2017).

The PPMI database follows specific eligibility criteria for *de novo* PD patients and healthy subjects. PPMI also provides metadata, including demographic and clinical information. Based on the reported Movement Disorders Society - Unified Parkinson’s Disease Rating Scale (MDS-UPDRS) part III/IV scores, we classified the selected PD patients. Those who had dyskinesia on examination (part III) and/or history (part IV) in any of their visits were classified as dyskinetic. Those who did not experience dyskinesia for at least 3 years since diagnosis were labeled as non-dyskinetic. For the baseline structural MRI analysis, we included 120 dyskinetics, 104 non-dyskinetics, and 100 healthy controls.

#### Resting-state functional MRI data

We selected a subset of 86 PD patients and 20 healthy subjects from the PPMI cohort who had both resting-state fMRI (rs-fMRI) and corresponding anatomical MPRAGE scans from any of their scheduled visits prior to LID onset. Of the 86 PD patients, 44 were classified as dyskinetics and 42 as non-dyskinetics. To make the group sizes comparable for analysis, we added 24 age and sex-matched healthy controls from the Cam-CAN database (https://www.cam-can.org) with similar imaging parameters (see supplementary material, Table S2) to the 20 age-and sex-matched healthy controls from the PPMI cohort. For all subjects, quality checks were performed with the CONN toolbox (Whitfield-Gabrieli S and Nieto-Castanon A 2012), and subjects with excessive head motion were excluded (see Data preprocessing section). In the final rs-fMRI dataset, we included data from 36 dyskinetics, 34 non-dyskinetics, and 32 healthy subjects. Of note, at the time the fMRI scans were obtained, some patients in both groups were already receiving levodopa treatment (see supplementary material, Table S5).

### Demographics and Clinical Characteristics

We extracted the following demographic and clinical data from the PPMI: age, sex, handedness, Hoehn & Yahr (H&Y) disease stage, disease duration, dyskinesia-free period, MDS-UPDRS scores (including the motor scores for the left and right sides separately), levodopa equivalent daily dose (LEDD), and scores on the Montreal Cognitive Assessment (MoCA) test and Geriatric Depression Scale (GDS) (see Methodology in supplementary material).

### Data preprocessing

#### Volumetric analysis

We used the standard preprocessing pipeline of the Freesurfer software version 7 (http://surfer.nmr.mgh.harvard.edu/). Briefly, we used the automated subcortical volume-based stream (Fischl et al., 2002) and the cortical surface-based stream (Dale et al., 1999, Fischl et al., 1999) to obtain the individual subcortical volumes, cortical volumes and cortical thickness values, respectively. We then performed visual quality checks of the preprocessed data. To test our hypothesis, the cortical thickness and volume of four cortical regions of interest (ROIs) including the precentral, postcentral, paracentral and frontal middle gyrus, and the volume of four subcortical ROIs including the caudate, putamen, globus pallidus, and thalamus were obtained.

#### Vertex-based surface analysis

We used the automated FMRIB laboratory’s Integrated Registration & Segmentation Tool (FSL FIRST Version 6.0.5) to segment the raw T1w image into subcortical structures (Patenaude et al., 2011). We used the segmentation data of the caudate, putamen, globus pallidus and thalamus for the subsequent steps. We used the FSL’s vertex analysis pipeline, which creates deformable meshes of the subcortical structures and fits them to a model derived from 336 subjects. The model contains manually segmented data of subcortical structures spanning both healthy controls and pathological brains. After registration and segmentation, we checked for errors using the ‘*first_roi_slicedir’* command before proceeding with shape analysis. We also extracted scalar projection values for each subject using the FSL Utilis Toolbox. Specifically, we used the *‘fslmaths’* command on the output of the vertex analysis to derive the scalar projection values. The scalar projection values depict mean perpendicular displacements of the vertices over the complete surface of the subcortical structure in comparison with the model.

#### Resting-state fMRI analysis

We used the *’dcm2niix’* toolbox of the MRIcroGL 64-bit software to convert the rs-fMRI scans from DICOM to NIFTI format. We used the CONN toolbox version 20.b (Whitfield-Gabrieli S and Nieto-Castanon A 2012) for all subsequent analyses. While preprocessing, we removed the initial four scans to eliminate magnetic equilibration effects. We then registered all scans to the initial scan of each session using b-spline interpolation and aligned them to the (0,0,0) coordinates across all sessions. We identified the outlier scans using ART (Artifact Detection Tools). Subsequently, we segmented functional and structural data into gray matter, white matter and cerebrospinal fluid, and normalized them to the standard Montreal Neurological Institute (MNI) brain template. We then applied a spatial smoothing kernel with an 8-mm full-width at half-maximum. We denoised the data by conducting linear detrending and despiking before regression. Our denoising approach followed the component-based noise reduction method described by Chai et al., 2012. During denoising, we implemented a bandpass filter to restrict the data to a frequency range of 0.008 < f < 0.09 Hz, capturing fluctuations in the blood oxygenation level-dependent (BOLD) signal, which predominantly occurs within this frequency range during resting states. We excluded subjects with invalid scans, mean head motion (i.e., 0.33 mm) deviating by more than three standard deviations from the mean or maximum head motion exceeding 3mm. Additionally, we computed the global correlation index (GCOR), which represents the average correlation coefficient between each pair of voxels across the entire brain for each subject. Here, GCOR serves as a quality control metric by calculating the distribution of the voxel-wise correlations after denoising.

### Statistical analysis

#### Overall analysis plan

First, we tested the normality of the distributions of all data. In cases of non-normal distribution, we performed non-parametric tests when applicable or excluded outliers to restore normality for the general linear model (GLM). For imaging analyses, we used the hypothesis-driven ROIs as dependent variables. We then conducted group analyses followed by planned pairwise comparisons. We added variables that exhibited significant differences among the groups as a covariates to the GLM.

#### Demographic and clinical data

We compared the cohort characteristics using the SPSS software version 29 (IBM Corporation, Armonk, NY, US). We tested the normality of the distribution using the Shapiro-Wilk test. If the distribution was non-normal, we used a non-parametric Mann-Whitney test for group comparisons. For normally distributed continuous variables, we used unpaired t-test. We used the Chi-square test to compare categorical variables.

#### Volumetric and cortical thickness data analysis

We extracted the cortical and subcortical volumes from the output files of the FreeSurfer automated segmentation pipeline and normalized them to the estimated total intracranial volume (eTIV). For the cortical thickness analysis, we obtained the thickness measures for parcellated cortical regions based on the Desikan-Killiany atlas from the FreeSurfer output files (see supplementary material, Table S3 & S4). The following analytical plan was applied to all volumetric and cortical thickness data: The left and right hemispheric measurements for each ROI were averaged so as to reduce the number of variables for multiple comparisons. We identified univariate outliers from group-wise box plots of each variable. Outliers exceeding ±2.7 S.D. from mean (determined by the IQR method (Mishra P et al., 2019)) were excluded from further analysis. We then conducted separate univariate one-way ANCOVAs to examine the main effect of the group (corrected p < 0.0125 for four cortical and subcortical ROIs). Significant differences were further examined by post-hoc pairwise comparisons using t-tests (Tukey’s correction at p < 0.05). We also conducted separate GLM analysis comparing dyskinetic and non-dyskinetic PD subjects using age, disease duration and dyskinesia-free period as covariates to account for the potential confounding effect of these variables. Significant differences were followed by post-hoc pairwise comparisons using t-tests (Tukey’s correction at p < 0.05). We used SPSS version 29 for our statistical analyses.

#### Vertex-based surface analysis

For the shape analysis, we used the *‘Randomise’* function of FSL to compare the groups. We performed an F-test across the three groups, followed by post hoc pairwise group comparisons. We performed 15,000 permutations for F-tests and 10,000 permutations per left and right subcortical structure for between-group comparisons with unpaired t-tests. We set contrasts to check for both “growth” and “atrophy” of the subcortical structures. We used age as a nuisance covariate. We used the Threshold Free Cluster Enhancement (TFCE) method for cluster definition. The results generated by randomizing were then corrected for multiple comparisons (Family Wise Error was set at p < 0.05). We used the cluster tool on the data corrected for multiple comparisons to report the cluster size and related cluster information such as extent, intensity, and associated t-statistics.

For the scalar projections analysis, we had eight dependent variables (four subcortical ROIs with their respective left and right hemispheric sides, corrected p < 0.006). The distribution of the data was analyzed using boxplots and histograms. Outliers were excluded from further univariate ANOVAs in SPSS to retain normality in the data. Significant group differences were then examined with Tukey’s post-hoc tests.

#### Resting-state fMRI analysis

We compared the GCOR values across groups using the GLM after denoising the BOLD signal. We then performed ROI-to-ROI connectivity (RRC) analyses using the CONN toolbox. We selected eight ROIs on each hemisphere defined using the FSL Harvard-Oxford atlas: Middle frontal gyrus (MFG), precentral gyrus (PreCG), postcentral gyrus (PostCG), supplementary motor area (SMA), inferior frontal gyrus (IFG), caudate, putamen, pallidum, and thalamus. For each subject, we extracted the average BOLD signal time courses from these ROIs and correlated them with each other. The ‘r’ values corresponded to the FC strength between ROI pairs. We Fisher z-transformed the ‘r’ values and obtained group-level FC maps for statistical analyses. We performed F-test for the three groups using multivariate parametric statistics with a connection threshold of p-uncorrected < 0.01 and a cluster threshold of p-FDR corrected < 0.05. Significant connections in the F-test were followed by post hoc pairwise group comparisons using the same multivariate parametric statistics and significance thresholds.

## Results

### Demographic and clinical characteristics

For the structural MRI analysis, the group sizes were comparable. There was no significant difference in sex, but a significant difference in age. Non-dyskinetic subjects were significantly older compared to dyskinetic and healthy subjects. Most subjects in each group were White. The PD subgroups showed significant differences in disease duration and, as expected, in dyskinesia-free period, but no significant difference in MDS-UPDRS I (non-motor experiences of daily living), MDS-UPDRS III (motor examination) and total MDS-UPDRS scores (combining parts I, II and III). We also compared the MDS-UPDRS III score separately for the left and right sides of the body (see Methodology in supplementary material). The scores were not significantly different between dyskinetics and non-dyskinetics, suggesting that the motor severity on either side of the body was comparable between the two patient subgroups. However, they differed in MDS-UPDRS II (motor experiences of daily living) scores. The MoCA scores obtained in the screening visit were also significantly different among the groups. Healthy subjects had significantly higher MoCA scores than non-dyskinetics. The descriptive statistics for demographic and clinical data are summarized in Table 1.

**Table 1:** Demographic and clinical characteristics of subjects in the structural MRI analysis.

|  | sMRI analysis participants |  |  | Corrected p-value |
| --- | --- | --- | --- | --- |
|  | Dyskinetics (n=120) | Non-Dyskinetics (n=104) | Healthy Subjects (n=100) |  |
| Age (y) | 59.50 ± 9.44 | 63.30 ± 8.37 | 59.8 ± 11.8 | **0.009<sup>a</sup>**, **1<sup>b</sup>**, **0.014<sup>c</sup>**, **0.040<sup>d</sup>** |
| Gender (M/F) | 73/47 | 65/39 | 65/35 | 0.816<sup>f</sup> |
| PD Duration at Baseline (y) | 0.11 ± 0.12 | 0.71 ± 0.76 | NA | **0.000<sup>e</sup>** |
| Dyskinesia free period (y) | 5.57 ± 2.53 | 7.47 ± 3.90 | NA | **0.000<sup>e</sup>** |
| Race & Ethnicity |  |  |  |  |
| Hispanic or Latino | 9 | 0 | 4 | - |
| Asian | 3 | 2 | 0 | - |
| Black/African American | 1 | 2 | 6 | - |
| Indian/Alaska Native | 1 | 1 | 3 | - |
| Race unspecified | 0 | 0 | 1 | - |
| White | 114 | 101 | 92 | - |
| H&Y | 1.65 ± 0.57 | 1.60 ± 0.49 | NA | 0.465<sup>e</sup> |
| MDS-UPDRS Total | 34.53 ± 14.20 | 31.7 ± 12.30 | NA | 0.118<sup>e</sup> |
| MDS-UPDRS I | 4.96 ± 3.27 | 4.19 ± 3.24 | NA | 0.080<sup>e</sup> |
| MDS-UPDRS II | 6.92 ± 4.54 | 4.82 ± 3.60 | NA | **0.000<sup>e</sup>** |
| MDS-UPDRS III | 20.91 ± 8.84 | 21.70 ± 9.01 | NA | 0.475<sup>e</sup> |
| MDS-UPDRS (R) | 6.14 ± 4.56 | 7.28 ± 4.54 | NA | 0.061<sup>e</sup> |
| MDS-UPDRS (L) | 6.65 ± 5.07 | 5.80 ± 5.01 | NA | 0.213<sup>e</sup> |
| MoCA (Screening visit) | 27.58 ± 2.24 | 27.25 ± 2.13 | 28.30 ± 1.24 | **0.004<sup>g</sup>**, 0.453<sup>h</sup>, 0.136<sup>i</sup>, **0.002<sup>j</sup>** |

M/F: Male/Female, MDS-UPDRS: Movement Disorders Society Unified Parkinson's Disease Rating Scale (L/R: Left- and right-sided motor exam scores), MoCA: Montreal Cognitive Assessment Score, PD: Parkinson's Disease, y: years. Numeric variables are reported as mean ± standard deviation. a: ANOVA, b: unpaired t-test (dyskinetics vs healthy controls), c: unpaired t-test (dyskinetics vs non-dyskinetics), d: unpaired t-test (healthy controls vs non-dyskinetics), f: Chi-square test, g: Kruskal-Wallis test, h: Dunn's test (dyskinetics vs non-dyskinetics), i: Dunn's test (dyskinetics vs healthy controls), j: Dunn's test (non-dyskinetics vs healthy controls).

In the fMRI dataset, there was no significant difference in age and sex, and the group sizes were comparable. Dyskinetic and non-dyskinetic subjects showed no significant difference in disease duration, MDS-UPDRS scores (I, II, III, & Total), Hoehn & Yahr disease stage, MoCA and GDS scores, or LEDD. The MDS-UPDRS III scores separately for the left and right sides of the body (see Methodology in supplementary material) were not significantly different between dyskinetics and non-dyskinetics, suggesting that the motor severity on either side of the body was comparable between the two patient subgroups. As expected, the PD subgroups significantly differed in the dyskinesia-free period. Table 2 summarizes the demographic and clinical characterisitcs of the subjects included in the fMRI dataset.

**Table 2:** Demographic and clinical characteristics of subjects included in the rs-fMRI analysis.

|  | fMRI analysis participants |  |  | Corrected p-value |
| --- | --- | --- | --- | --- |
|  | Dyskinetics (n=36) | Non- Dyskinetics (n=34) | Healthy Controls (n=32) |  |
| Age (y) | 60.01 ± 9.02 | 64.63 ± 10.01 | 61.34 ± 9.80 | 0.081<sup>a</sup> |
| Sex (M/F) | 22/14 | 23/9 | 22/12 | 0.639<sup>b</sup> |
| PD Duration (y) | 2.08 ± 1.51 | 2.50 ± 2.74 | NA | 0.971<sup>d</sup> |
| Dyskinesia free period (y) | 5.02 ± 2.31 | 7.94 ± 2.62 | NA | **0.000<sup>c</sup>** |
| MDS-UPDRS I | 5.14 ± 3.52 | 4.76 ± 3.12 | NA | 0.637<sup>d</sup> |
| MDS-UPDRS II | 7.38 ± 3.75 | 6.11 ± 4.57 | NA | 0.088<sup>d</sup> |
| MDS-UPDRS III | 22.38 ± 11.46 | 21.64 ± 11.04 | NA | 0.783<sup>c</sup> |
| MDS-UPDRS Total | 34.80 ± 14.33 | 32.91 ± 15.32 | NA | 0.594<sup>c</sup> |
| MoCA | 20.11 ± 12.07 | 18.15 ± 12.24 | NA | 0.308<sup>d</sup> |
| GDS | 2.55 ± 2.62 | 1.67 ± 1.87 | NA | 0.175<sup>d</sup> |
| LEDD | 485.60 ± 477.66 | 513.30 ± 630.91 | NA | 0.656<sup>d</sup> |
| H&Y | 1.83 ± 0.44 | 1.76 ± 0.43 | NA | 0.588<sup>d</sup> |
| MDS-UPDRS (L) | 7.11 ± 5.30 | 5.45 ± 4.91 | NA | 0.199<sup>d</sup> |
| MDS-UPDRS (R) | 6.66 ± 4.22 | 7.06 ± 4.16 | NA | 0.697<sup>c</sup> |

GDS: Geriatric Depression Scale, H&Y: Hoehn & Yahr Scale, LEDD: L-dopa equivalent daily dose, M/F: Male/Female, MoCA: Montreal Cognitive Assessment, MDS-UPDRS: Movement Disorders Society Unified Parkinson's Disease Rating Scale, PD: Parkinson's Disease (L/R: Left- and right-sided motor exam scores), y: years. Numeric variables are reported as mean ± standard deviation. a: Kruskal-Wallis test, b: Chi-square test, c: Unpaired t-test (dyskinetic vs non-dyskinetic), d: Mann-Whitney test (dyskinetic vs non-dyskinetic).

### Volumetric analysis

#### Cortical volume

The final cohort sizes after removing the outliers were as follows: dyskinetics (n=110), non-dyskinetics (n=95) and healthy controls (n=92). The distribution of the data was normal (see supplementary material, Figure S1). The ANCOVA with age as a covariate did not show any significant volumetric difference among the groups. The comparison between dyskinetics and non-dyskinetics with age, disease duration and dyskinesia-free period as covariates also did not show significant volumetric differences.

#### Cortical thickness

The final cohort sizes after removing the outliers were as follows: dyskinetics (n=113), non-dyskinetics (n=98) and healthy controls (n=93). The distribution of the data was normal (see supplementary material, Figure S3). The ANCOVA did not show any significant difference in cortical thicknesses among the groups. However, the comparison between dyskinetics and non-dyskinetics with age, disease duration and dyskinesia-free period as covariates showed significant differences. The dyskinetics had a significantly higher thickness in the precentral (F (1, 219) = 7.045, p =0.009), paracentral (F (1, 219) = 6.826, p=0.010), middle frontal (F (1, 219) = 9.837, p = 0.002) and postcentral (F (1, 219) = 8.88, p = 0.003) regions than non-dyskinetics (Figure 2).

**Figure 2:**
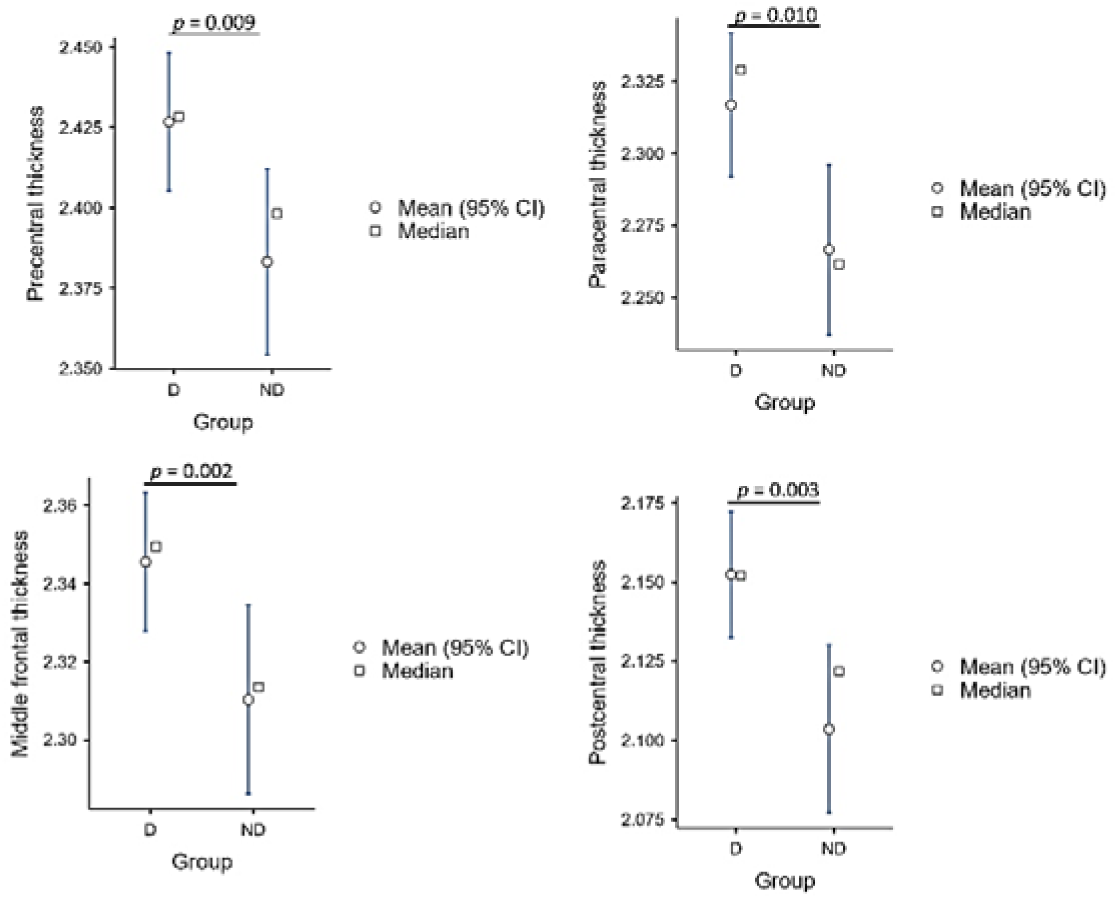
Cortical thicknesses in dyskinetic and non-dyskinetic PD patients. Mean (with 95% confidence interval) and median plots showing precentral, paracentral, middle frontal and postcentral gyrus thickness. D: Dyskinetics, ND: Non-dyskinetics.

#### Subcortical volume

The final cohort sizes after removing the outliers were as follows: dyskinetics (n=114), non-dyskinetics (n=100) and healthy controls (n=97). The distribution of the data was normal (see supplementary material, Figure S5). The ANCOVA did not show any significant differences in the subcortical volumes among the groups. The subcortical volumes also did not differ significantly between dyskinetics and non-dyskinetics after controlling for age, disease duration and dyskinesia-free period. Nonetheless, we observed a significant effect of age on subcortical volumes (see supplementary material, Figure S13).

### Vertex-based surface analysis

The ANOVA demonstrated significant surface clusters in the left pallidum, right pallidum, left caudate and left putamen (p < 0.05). The post-hoc pairwise comparisons showed significant clusters in the left caudate and left pallidum between dyskinetics and non-dyskinetics and a significant cluster in the left pallidum between dyskinetics and healthy controls. No significant clusters were found between non-dyskinetics and healthy controls (Table 3, Figure 3). We also compared dyskinetics and non-dyskinetics separately after controlling for ‘age’ and ‘dyskinesia-free period’. We found significant clusters in the left caudate at the exact location shown before (see supplementary material, Table S7).

**Figure 3:**
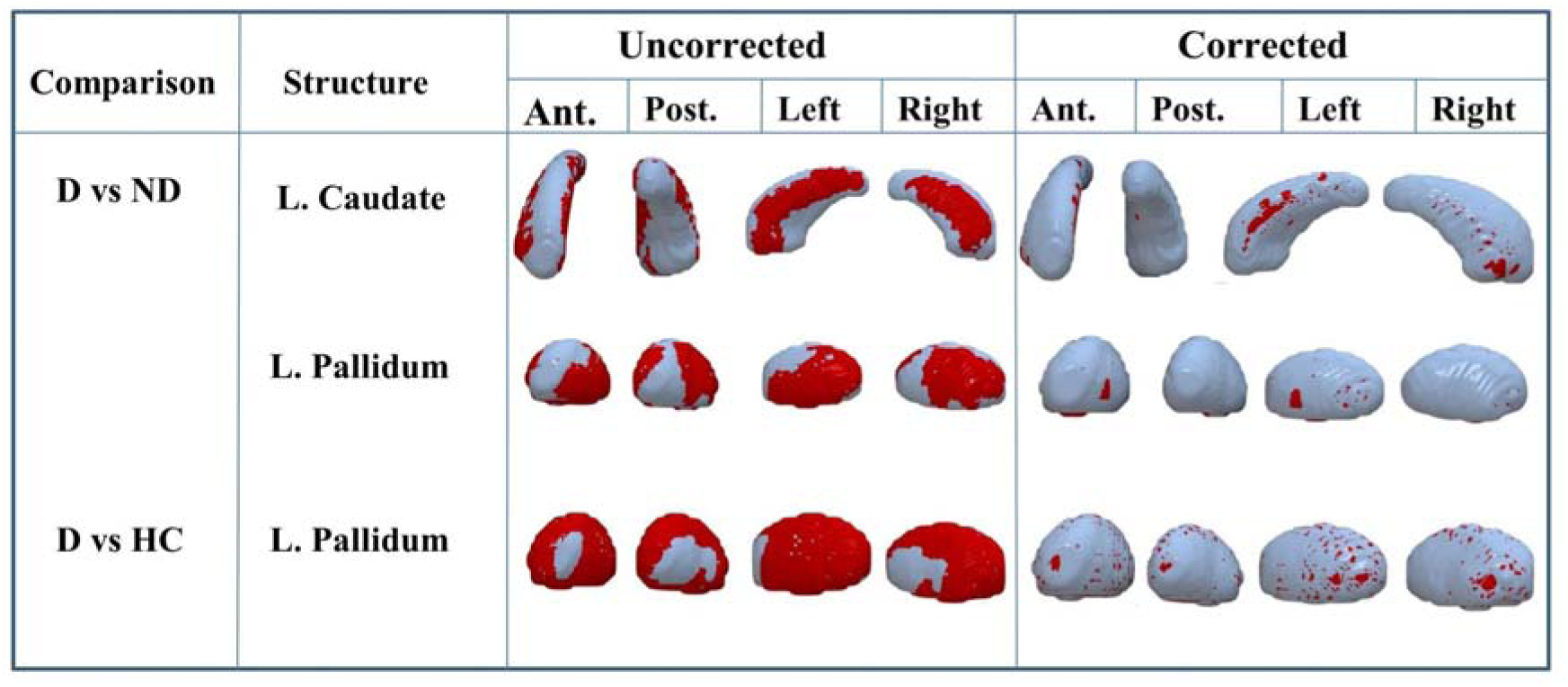
Maps of surface clusters that showed significant between-group differences. The red clusters are shown from anterior, posterior, left, and right views at uncorrected and corrected (FWE, p<0.05) thresholds. D: Dyskinetics, HC: Healthy controls, ND: Non-dyskinetics.

**Table 3:** Basal ganglia vertex analysis results.

| Groups | Cluster no. | Vertex analysis |  | X | Y | Z |
| --- | --- | --- | --- | --- | --- | --- |
|  |  | Size | Max T-stat |  |  |  |
| **D > ND** |  |  |  |  |  |  |
| Left caudate | 1 | 377 | 3.33 | 99 | 132 | 84 |
| Left Caudate | 2 | 47 | 3.27 | 109 | 136 | 86 |
| Left Pallidum | 3 | 17 | 2.9 | 104 | 123 | 72 |
| Left Pallidum | 4 | 10 | 2.8 | 109 | 117 | 73 |
| **D > HC** |  |  |  |  |  |  |
| Left Pallidum | 5 | 420 | 2.9 | 108 | 121 | 76 |

### Scalar projection values

The final cohort sizes after removing the outliers were as follows: dyskinetics (n=115), non-dyskinetics (n=93) and healthy controls (n=92). The ANOVA showed significant differences in the left pallidum (F (2,296) =5.00, p =0.007), right pallidum (F (2,296) = 5.02, p =0.007), left caudate (F (2,296) =3.64, p =0.027), right caudate (F (2,296) =3.51, p =0.031), and left putamen (F (2,296) =3.27, p =0.039). Only the left and right pallidum showed a strong trend towards significance (approaching p_corr._=0.006) after correction for multiple comparisons. This trend showed a net-outward deformation in dyskinetics compared to healthy controls in the pallidum (Figure 4). However, the dyskinetics did not differ significantly from the non-dyskinetics after controlling for age, disease duration and dyskinesia-free period.

**Figure 4.**
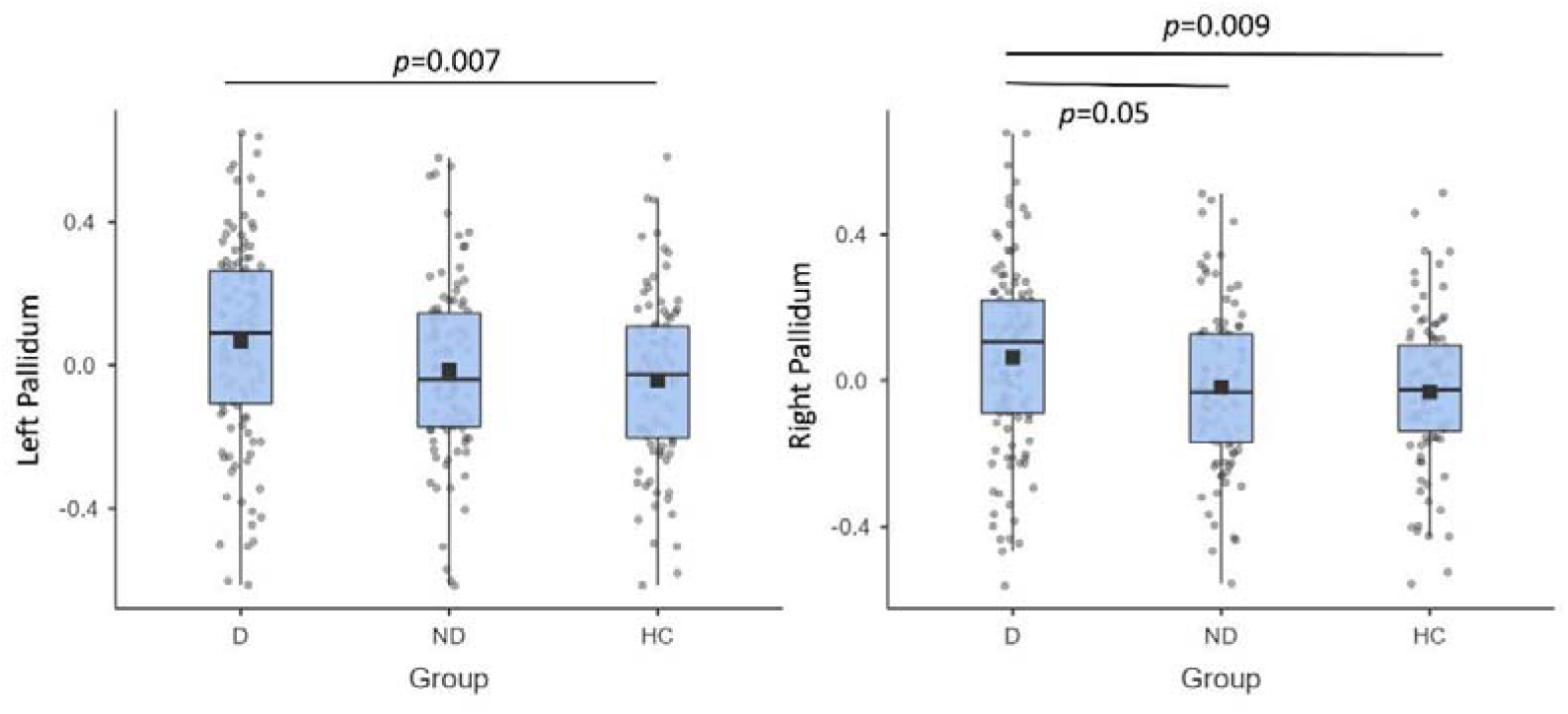
Scalar projection values. The box plots show the distribution of the scalar projection values of the left and right pallium for three groups. D: Dyskinetics, HC: Healthy controls, ND: Non-dyskinetics.

### Resting-state fMRI ROI-to-ROI connectivity analysis

There was a statistically non-significant, yet substantial age gap between dyskinetics and non-dyskinetics. Hence, we included age as a covariate in the FC analysis. Eight out of 18 ROIs including the left putamen (F(12,186) = 4.63, p-FDR < 0.001), right putamen (F(12,186) = 4.53, p-FDR < 0.001), left inferior frontal (F(12,186) = 3.56, p-FDR < 0.001), right inferior frontal (F(12,186) = 2.22, p-FDR = 0.027), left postcentral F(12,186) = 2.35, p-FDR = 0.019), right postcentral F(12,186) = 2.45, p-FDR = 0.016), left precentral F(12,186) = 2.89, p-FDR = 0.003), and right precentral F(12,186) = 2.90, p-FDR = 0.003) gyri showed significant connectivity differences. The significant pairwise connectivity differences of these ROIs with the other ROIs are shown in Table 4 and Figure 5A.

**Figure 5:**
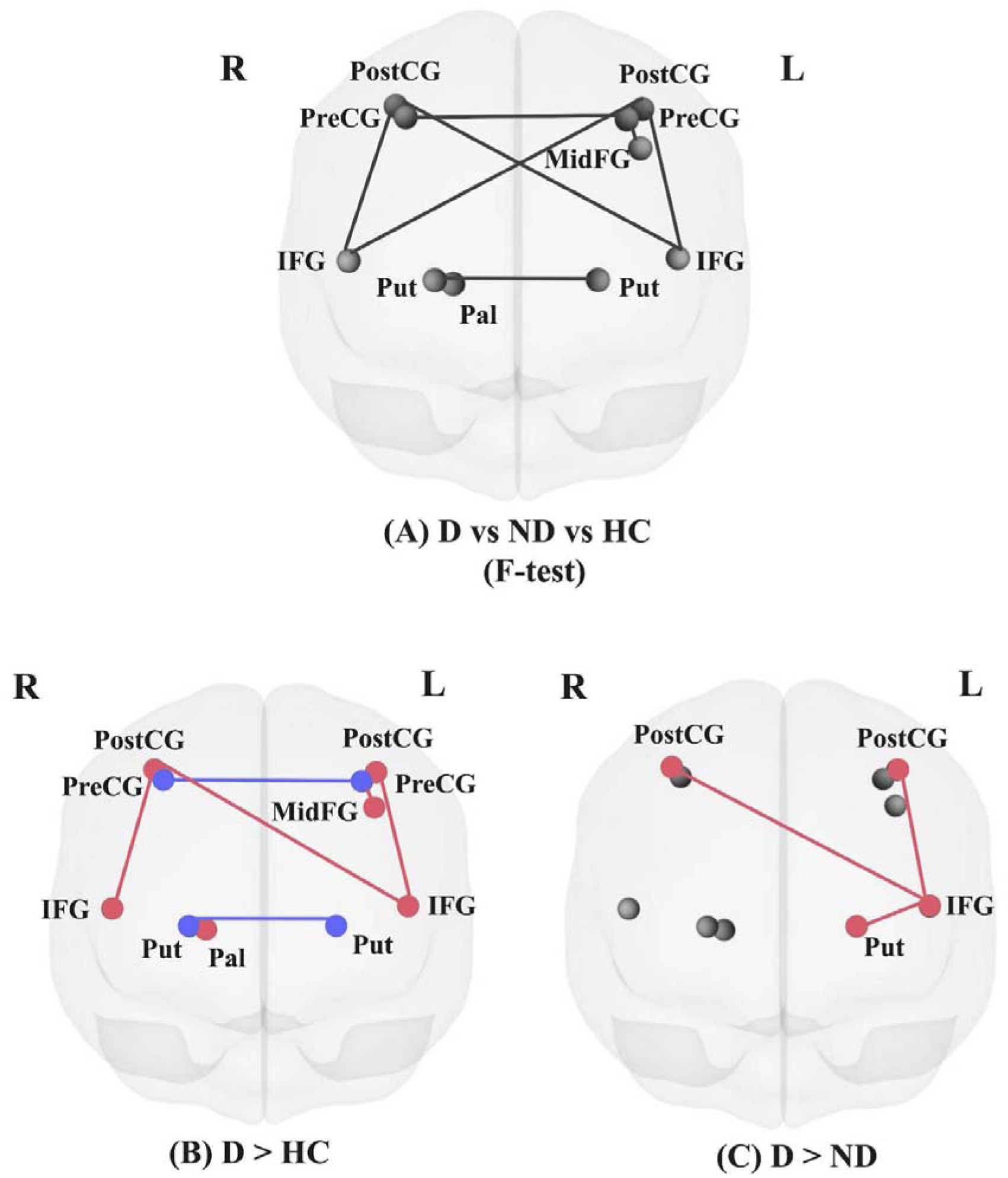
Pairwise group comparisons of ROI-to-ROI connectivity. (A) F-test results (D vs ND vs HC) followed by post hoc group comparisons: (B) D > HC, (C) D > ND. Cau: Caudate, D: Dyskinetics, HC: Healthy Controls, IFG: Inferior frontal gyrus, L: Left, MidFG: Middle frontal gyrus, ND: Non-Dyskinetics, Pal: Pallidum, PostCG: Postcentral gyrus, PreCG: Precentral gyrus, Put: Putamen, R: Right

**Table 4:** Pairwise connections that show group differences.

| **F-Statistics for ROI-to-ROI connectivity** |  |  |  |
| --- | --- | --- | --- |
| **Connections** |  | **F-Statistics** | **p-FDRcorr** |
| **ROI 1** | **ROI 2** |  |  |
| **Putamen l** | Putamen r | F (2, 98) = 18.35 | 0.000 |
| **Putamen l** | IFG l | F (2, 98) = 4.84 | 0.046 |
| **Putamen r** | Pallidum r | F (2, 98) = 5.87 | 0.033 |
| **IFG l** | PostCG l | F (2, 98) = 9.28 | 0.003 |
| **IFG l** | PostCG r | F (2, 98) = 6.95 | 0.012 |
| **IFG r** | PostCG l | F (2, 98) = 7.26 | 0.013 |
| **IFG r** | PostCG r | F (2, 98) = 6.93 | 0.013 |
| **PreCG l** | MidFG l | F (2, 98) = 7.23 | 0.019 |
| **PreCG l** | PreCG r | F (2, 98) = 5.62 | 0.041 |

ROI-to-ROI pairwise connections that show significant differences between the dyskinesics, non-dyskinesics, and healthy controls. IFG: Inferior frontal gyrus, l/r: left/right, MidFG: Middle frontal gyrus, PostCG: Postcentral gyrus, PreCG: Precentral gyrus.

We then performed post-hoc pairwise comparisons to determine the connectivity differences between the groups (Table 5). There was no significant difference between healthy controls and non-dyskinetics. However, dyskinetics compared to healthy controls showed significantly stronger connectivity among various cortical regions and between the putamen and pallidum, and significantly weaker connectivity between the left and right putamen (Figure 5B). Importantly, dyskinetics showed significantly stronger connectivity of the left inferior frontal gyrus with bilateral postcentral gyri and the left putamen compared to non-dyskinetics (Figure 5C). As the dyskinesia-free period was significantly different between the two PD subgroups, we also compared dyskinetics to non-dyskinetics using both dyskinesia-free period and age as covariates and found the same connectivity differences (see supplementary material, Figure S15).

**Table 5:** Pairwise group comparisons of ROI-to-ROI connectivity.

| **Pairwise comparisons for ROI-to-ROI connectivity** |  |  |  |
| --- | --- | --- | --- |
| **Connections** |  | **T-Statistics** | **p-FDRcorr** |
| **ROI 1** | **ROI 2** |  |  |
| **Dyskinetics > Non-Dyskinetics** |  |  |  |
| **IFG l** | PostCG l | T (67) = 3.40 | 0.019 |
| **IFG l** | PostCG r | T (67) = 3.02 | 0.030 |
| **IFG l** | Putamen l | T (67) = 2.75 | 0.032 |
| **Dyskinetics > Healthy Controls** |  |  |  |
| **Putamen l** | Putamen r | T (65) = -6.32 | 0.000 |
| **Putamen r** | Pallidum r | T (65) = 3.01 | 0.031 |
| **IFG l** | PostCG l | T (65) = 3.91 | 0.003 |
| **IFG l** | PostCG r | T (65) = 3.28 | 0.014 |
| **IFG r** | PostCG r | T (65) = 3.21 | 0.035 |
| **PreCG l** | MidFG l | T (65) = 3.84 | 0.004 |
| **PreCG l** | PreCG r | T (65) = -3.00 | 0.032 |

IFG: Inferior frontal gyrus, l/r: left/right, MidFG: Middle frontal Gyrus, PostCG: Postcentral gyrus, PreCG: Precentral gyrus.

## Discussion

To our knowledge, this is the first study to investigate differences in both structural and functional MRI from a large cohort of PD patients prior to the onset of LID. Specifically, our findings indicate significantly thicker frontal and sensorimotor cortices in dyskinetic compared to non-dyskinetic PD patients, even before initiating levodopa therapy. While subcortical structure volumes were similar between the groups, vertex-based shape analysis revealed notable surface inflation in the left caudate and left pallidum in the dyskinetic compared to the non-dykinetic group. Additionally, scalar projection values showed trends for pallidal shape differences in the dyskinetic group. Lastly, our fMRI analysis identified hyperconnectivity between the left IFG and the left putamen, as well as bilateral postcentral gyri, in dyskinetic patients before the clinical manifestation of LID.

### Volumetric changes

Cortical and subcortical volumetric differences between PD patients and healthy controls have been investigated extensively with conflicting results especially relatively early in the disease course (De Micco R et al., 2018, Pereira et al., 2012). As PD advances, patients tend to exhibit a decrease in the volume of specific brain regions more consistently, such as the putamen and frontal-parietal cortical areas (Lewis MM et al., 2012). Among the very few studies that investigated dyskinetic and non-dyskinetic PD patients separately, one study showed an increased IFG volume in dyskinetic PD patients (Cerasa A et al., 2011). But the PD patients in that study were already diagnosed with LID and were on medication for a minimum of six months. In our study, we did not see significant differences in the cortical or subcortical volumes between the PD groups suggesting that baseline volumetric characteristics do not distinguish between dyskinetic and non-dyskinetic PD patients at an early (*de novo*) stage.

### Cortical thickness changes

Past studies suggest widespread cortical thinning in PD patients compared to healthy subjects (De Micco R et al., 2018) at different stages of PD progression (Wilson H et al., 2019), especially in those with cognitive decline (Cicero CE et al., 2022). Specific to LID, increased thickness of the right inferior frontal sulcus in PD patients with dyskinesia compared to those without has been reported (Cerasa A et al., 2013). Similarly, our results showed an increased thickness of the precentral, paracentral, middle frontal and postcentral gyri in dyskinetics compared to non-dyskinetics. A recent study suggests that the brain-derived neurotrophic factor gene rs6265 single nucleotide polymorphism may be responsible for an increased cortical thickness in the left postcentral gyrus in PD patients with LID and is likely to be linked to the development of dyskinesia (Sun HM et al., 2022). These findings collectively indicate that certain PD patients may possess a genetic predisposition to dyskinesia, potentially driven by morphological alterations in the sensorimotor cortex.

### Basal ganglia shape changes

Consistent and significant basal ganglia shape changes in PD patients compared to healthy subjects have been demonstrated across two independent datasets. These changes were characterized by atrophy (net-inward deformation/shrinkage) in the right putamen and bilateral pallidum, and by signs of both atrophy and hypertrophy in the left thalamus and bilateral caudate in the PD group (Garg et al., 2015). Using the large ENIGMA-PD consortium database, a recent study showed basal ganglia shape changes in PD patients at various diesase stages compared to matched controls. These changes were characterized by progressive thinning of these structures across disease stages starting with the putamen followed by the caudate, nucleus accumbens, and globus pallidus. Interestingly, earlier in the disease course, subregions of the thalamus were found to be thicker (Laansma et al., 2024). Importantly, another recent study also showed local shape changes in the left putamen and left caudate in dyskinetic compared to non-dyskinetic PD patients (Kim et al., 2020) suggesting that these striatal shape changes have the potential to be used as a surface-based biomarker related to LID severity. We also found subtle but significant shape changes, i.e., clusters of surface growth in the left caudate and left pallidum in the dyskinetic patient group. Additionally, the scalar projection values showed a trend for global surface growth in bilateral pallidum in dyskinetics compared to healthy controls and in the right pallidum in dyskinetic compared to non-dyskinetics. This is in contrast with the results of a more recent study by Youn et al., 2022 that showed atrophy in right globus pallidus internus in dyskinetic compared to non-dyskinetic PD patients. However, in their study, the dyskinetic and non-dyskinetic groups differed in age, disease duration, motor severity, and LEDD. Animal models of PD further corroborate the role of the morphological changes in the globus pallidus in LID development. For example, hypertrophy in the medial globus pallidus was found to be associated with LID in 6-OHDA lesioned rats (Tomiyama et al., 2004). Moreover, younger rats with 6-OHDA lesions that were treated with levodopa were found to express more enhanced LID-like behavior and larger medial globus pallidus volumes compared to older rats (Nishijima et al., 2021). The globus pallidus internus is a well-established neurosurgical target for deep brain stimulation to treat LID in PD patients (Deuschl G. et al., 2006, Obeso, J. A et al., 2001). In light of our findings and previous reports, we think that presurgical planning might benefit from considering the morphological changes in this structure as revealed by surface analysis for better spatial accuracy in lead placement.

### Resting-state FC changes

Hypoconnectivity in the basal ganglia circuitry in PD patients compared to healthy subjects is well-established (Hacker CD et al., 2012, Szewczyk-Krolikowski K et al., 2014, Herz et al., 2014). Our findings also corroborate this connectivity pattern in dyskinetic PD patients, particularly hypoconnectivity between bilateral putamen. Additionally, we also report hyperconnectivity between the right putamen and pallidum. Dyskinetic patients also showed significant hyperconnectivity in the cortical circuitry compared to non-dyskinetics and healthy controls. Specifically, we found stronger connectivity of the left IFG with left putamen and bilateral somatosensory cortices in dyskinetic PD patients. Consistent with our findings, a more recent study showed an increased FC of the left IFG with the motor cerebellum (lobule VIIIb) in PD patients who developed LID within 5 years compared to those who did not (Yoo HS et al., 2019). Desynchronized FC of the IFG and pre-supplementary motor area during medication “on” state in dyskinetic compared to non-dyskinetic PD patients has also been shown suggesting uncoordinated inhibitory control over motor circuits as a mechanism underlying LID (Gan et al., 2020). On the contrary, reduced activity of the right IFG in a motor task (Cerasa et al., 2012) and a weaker resting-state FC between the right IFG and primary motor cortex (Cerasa et al., 2015) in dyskinetic vs non-dyskinetic patients have also been demonstrated before. Neuromodulation studies have also targeted the IFG for management of dyskinesia. For example, a transcranial magnetic stimulation (TMS) study showed that continuous theta burst stimulation, an inhibitory protocol, applied over the right IFG decreased dyskinesia severity (Cerasa et al., 2015). A follow-up study (Ponzo et al., 2016) using dual-site TMS revealed that there was reduced cortico-cortical inhibitory control of IFG over the contralateral motor cortex that resulted in abnormal facilitation of the motor cortex in dyskinetics. Taken together, these results suggest that IFG may be a key region in LID pathophysiology and a good target for noninvasive neuromodulation for the management of dyskinesia in PD patients. Our findings further emphasize the importance of exploring the potential role of IFG inhibition over the sensory cortex in the pathophysiology of dyskinesias.

## Conclusion

The current study has revealed structural and functional changes in the sensorimotor cortical-basal ganglia circuitry of de novo PD patients who later developed LID. Vertex-based shape analysis identified subtle morphological changes in the striatum and globus pallidus that standard volumetric analyses failed to detect. Additionally, our findings point to interhemispheric basal ganglia functional dysconnectivity and abnormal cortical hyperconnectivity in dyskinetic patients. Previous reports indicate a direct cortico-cortical modulatory role for the IFG on the motor cortex activity which seems disrupted in dyskinetic patients. Our IFG-sensory cortex hyperconnectivity finding in the dyskinetic group is novel and suggests a similar role for the IFG on the sensory cortex.

The strengths of our study are the use of robust multimodal imaging methodology to probe the structure and function of the cortical-basal ganglia circuits and inclusion of relatively large, well-characterized and comparable de novo PD cohorts that allowed us to minimize the confounding effects of demographic and clinical differences. However, this study also has limitations. The data in the PPMI database were collected at different centers, and unlike the structural MRI scans that were acquired at baseline, the fMRI scans were collected at different time points within two years of enrollment (but before the onset of dyskinesia). An important next step would be the replication of our findings combined with electrophysiological studies in well-characterized prospective PD cohorts.

## CrediT authorship contribution statement

**Sakshi Shukla:** Data curation, Software, Formal analysis, Validation, Writing-Original draft, Writing-Review & Editing, Visualization. **Mantosh Patnaik:** Data curation, Software, Formal analysis, Visualization. **Aditya Kumar:** Data curation, Software, Formal analysis. **Sule Tinaz:** Conceptualization, Methodology, Resources, Writing-Original draft, Writing-Review & Editing, Supervision, Project Administration. **Nivethida Thirugnanasambandam:** Conceptualization, Methodology, Resources, Writing-Original draft, Writing-Review & Editing, Supervision, Project Administration.

## Acknowledgements

This work was supported by the DBT/Wellcome Trust India Alliance Fellowship [grant number: IA/CPHI/16/1/502624] awarded to NT. SS was partially supported by the Fulbright - Nehru doctoral research fellowship.

## Data Availability

Data is available from the PPMI database (www.ppmi-info.org/data).

