## Supplementary material for "Multimodal Neuroimaging Reveals Distinct Characteristics of Levodopa-Induced Dyskinesias in *de novo* Parkinson’s Disease Patients"

**Methodology**

**Demographic and Clinical Characteristics**

The information about the subjects’ age, sex, handedness, and Hoehn & Yahr Stage was provided under the PPMI Laboratory of Neuroimaging (LONI) platform > Download > Study data in the PPMI database. Based on the information provided, we calculated the following additional parameters for our study:

- **Total MDS-UPDRS score** - We combined the MDS-UPDRS score of Part I: Non-Motor aspects of experiences of daily living, Part II: Motor aspects of experiences of daily living, Part III: Motor examination and Part IV: Motor complications.
- **MDS-UPDRS score for the left side of the body –** A cumulative motor impairment score was calculated for the left side based on the following MDS-UPDRS part III items: 3.3c Rigidity in the left upper extremity, 3.3e Rigidity in the left lower extremity, 3.4b Finger tapping in the left hand, 3.5b Hand movements in the left hand, 3.6b Pronation-supination movements-left hand, 3.7b Toe tapping in the left foot, 3.8b Leg agility in the left leg, 3.15b Postural tremor in the left hand, 3.16b Kinetic tremor in the left hand, 3.17b Rest tremor amplitude in left upper extremity, 3.17d Rest tremor amplitude in left lower extremity.
- **MDS-UPDRS score for right side of the body -** A cumulative motor impairment score was calculated for the right side based on the following MDS-UPDRS part III: 3.3b Rigidity in right upper extremity, 3.3d Rigidity in right lower extremity, 3.4a Finger tapping in right hand, 3.5a Hand movements in right hand, 3.6a Pronation-supination movements-right hand, 3.7a Toe tapping in right foot, 3.8a Leg agility in right leg, 3.15a Postural tremor in right hand, 3.16a Kinetic tremor in right hand, 3.17a Rest tremor amplitude right upper extremity, 3.17c Rest tremor amplitude in right lower extremity.
- **Affected side –** We compared the MDS-UPDRS score for each subject's left and right sides of the body. The side with the higher score was considered as the ‘more affected side’ for the subject.
- **Levodopa Equivalent Daily Dose (LEDD)-** We calculated the levodopa equivalent daily doses (LEDD) according to the Tomlinson CL et al., 2010 as described in the PPMI instructions. The medication information was provided in the PPMI database under the Download > Study data > Medical History > LEDD Concomitant medication log. We used our customized Python script to calculate the LEDD values for all subjects with respect to the time window of their imaging sessions.
- **Disease duration-** The disease duration corresponded to the number of months/years between the patient’s PD diagnosis date and their imaging session date (based on the visit considered in the analysis).
- **Dyskinesia-free period- For Dyskinetics:** Any marking regarding dyskinesia in part III or IV of the MDS-UPDRS rating was considered a sign of dyskinesia development in the subject. The duration from the date of diagnosis to the earliest visit where dyskinesia appeared was regarded as ‘Dyskinesia-free period’ in dyskinetic PD patients. **For Non-dyskinetics:** Patients for whom there was no marking of dyskinesia in the database were considered non-dyskinetic, and the time from their PD diagnosis date to their last visit was considered ‘Non-dyskinetic phase’.
- **Geriatric Depression Scale (GDS) –** We used the shorter version of GDS with 15 items. The information was provided under Download > Study data > Non-motor assessments > Neurobehavioral tests. According to the guidelines, we added 1 point for each response of ‘No’ (0) to any of the following variables: GDSSATIS, GDSGSPIR, GDSHAPPY, GDSALIVE, GDSENERGY. And we added 1 point for each response of ‘Yes’ (1) to any of the following variables: GDSDROPD, GDSEMPTY, GDSBORED, GDSAFRAID, GDSHLPLS, GDSHOME, GDSMEMRY, GDSWRTLS, GDSHOPLS, GDSBETER. Subjects with a total GDS score greater or equal to 5 were listed as ‘Depressed’ while subjects with a score of less than 5 were considered ‘Not depressed’.
- **Montreal Cognitive Assessment (MoCA) –** The total MoCA score was already provided in the PPMI database under Download > Study data > Non-motor assessments > Neuropsychological tests. The MoCA score was unavailable for the baseline visit. Hence, we considered MoCA scores from the subject’s screening visits to be considered for the baseline data analysis.

We used customized python scripts to process and extract the above-mentioned parameters for all our subjects from the raw files together into a master sheet.

**Image acquisition and parameters**

The protocols for PPMI imaging data acquisition have been provided in the PPMI’s imaging technical operation manuals under research documents and standard operating procedures. All T1w structural and T2*GE EPI functional MRI scans were acquired in 3T scanners. Detailed information of the structural and functional scan sequences can be found in Table S1 and Table S2, respectively.

**Table S1: Description of the major PPMI structural MRI scan sequences.**

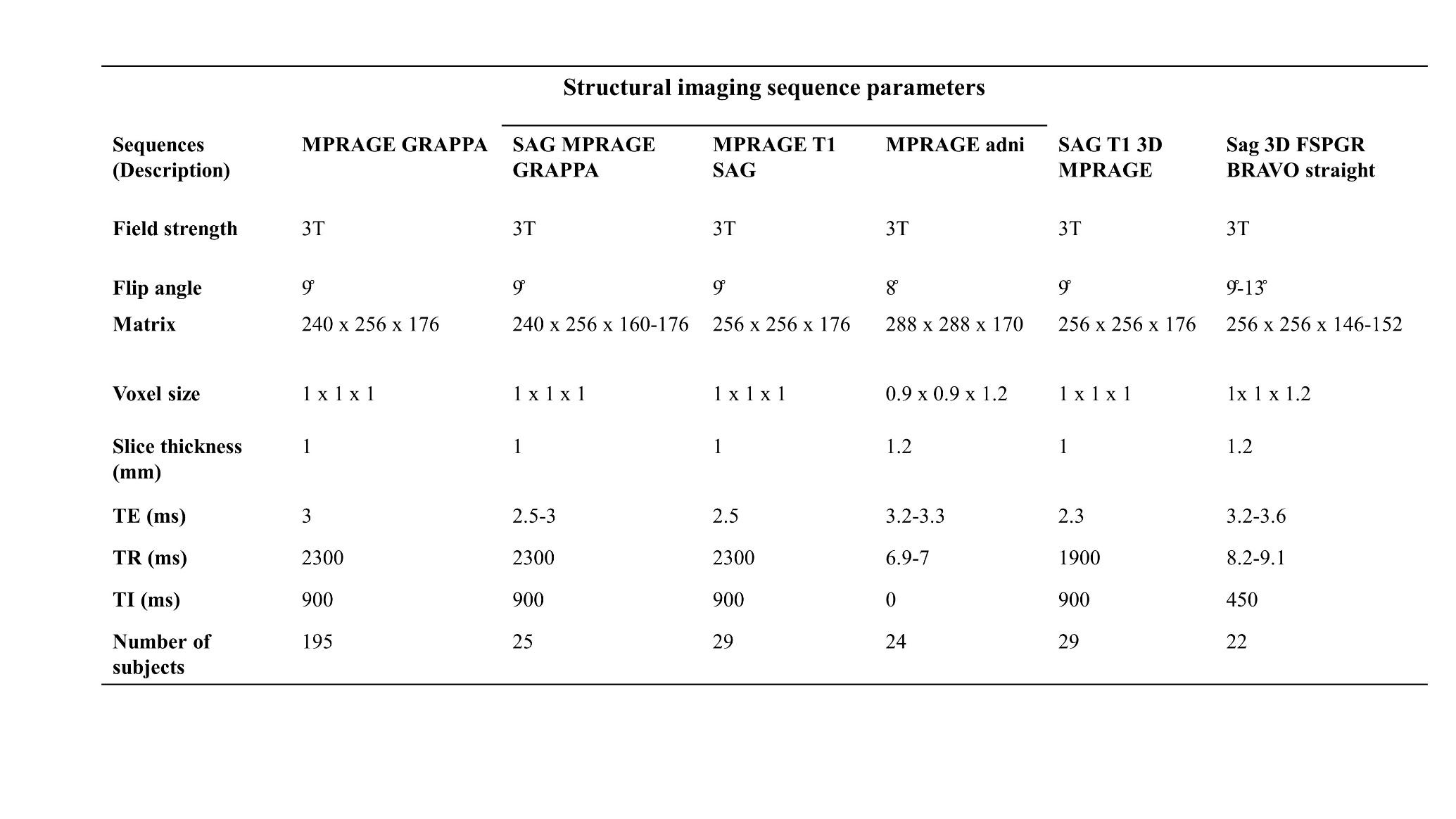

**Table S2: Description of the PPMI and Cam-CAN rs-fMRI scan sequences.**

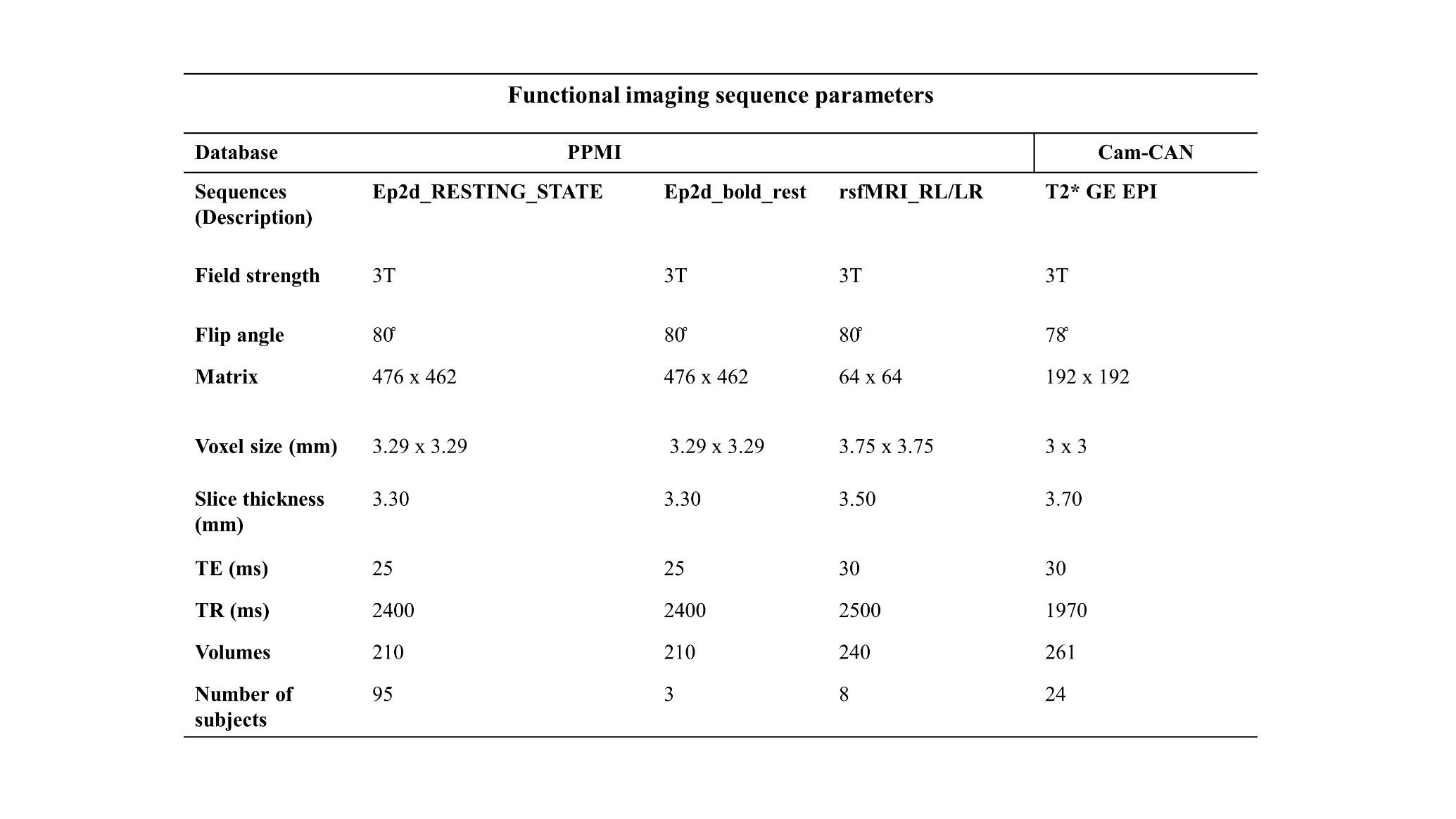

**Baseline structural MRI data**

In Freesurfer, the parcellation is based on ‘Destrieux’ cortical atlas (Destrieux C et al., 2010). In its parcellation technique, a cortical region is divided into subsequent gyral (visible cortex) and sulcal (hidden cortex) regions. Hence, we summed the subdivisions in the case of cortical volumes and averaged them in the case of cortical thicknesses. Unlike volumes, the thicknesses of the regions were not normalized to the average global thickness. After obtaining cortical volumes and thicknesses in each hemisphere, the values were averaged across the hemispheres **(**Tables S3 and S4).

**Table S3: Cortical volume calculations**

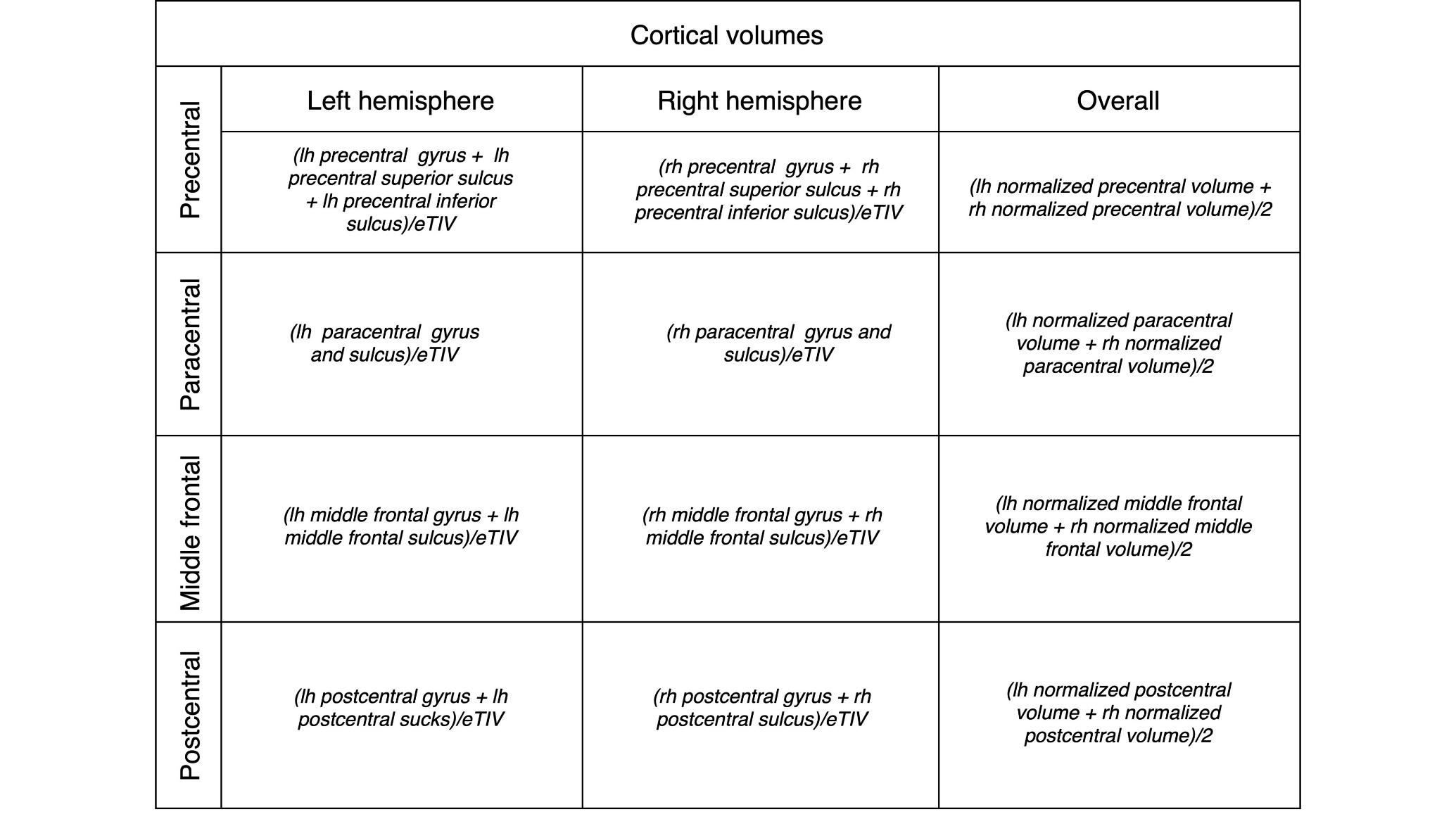

**Table S4: Cortical thickness calculations**

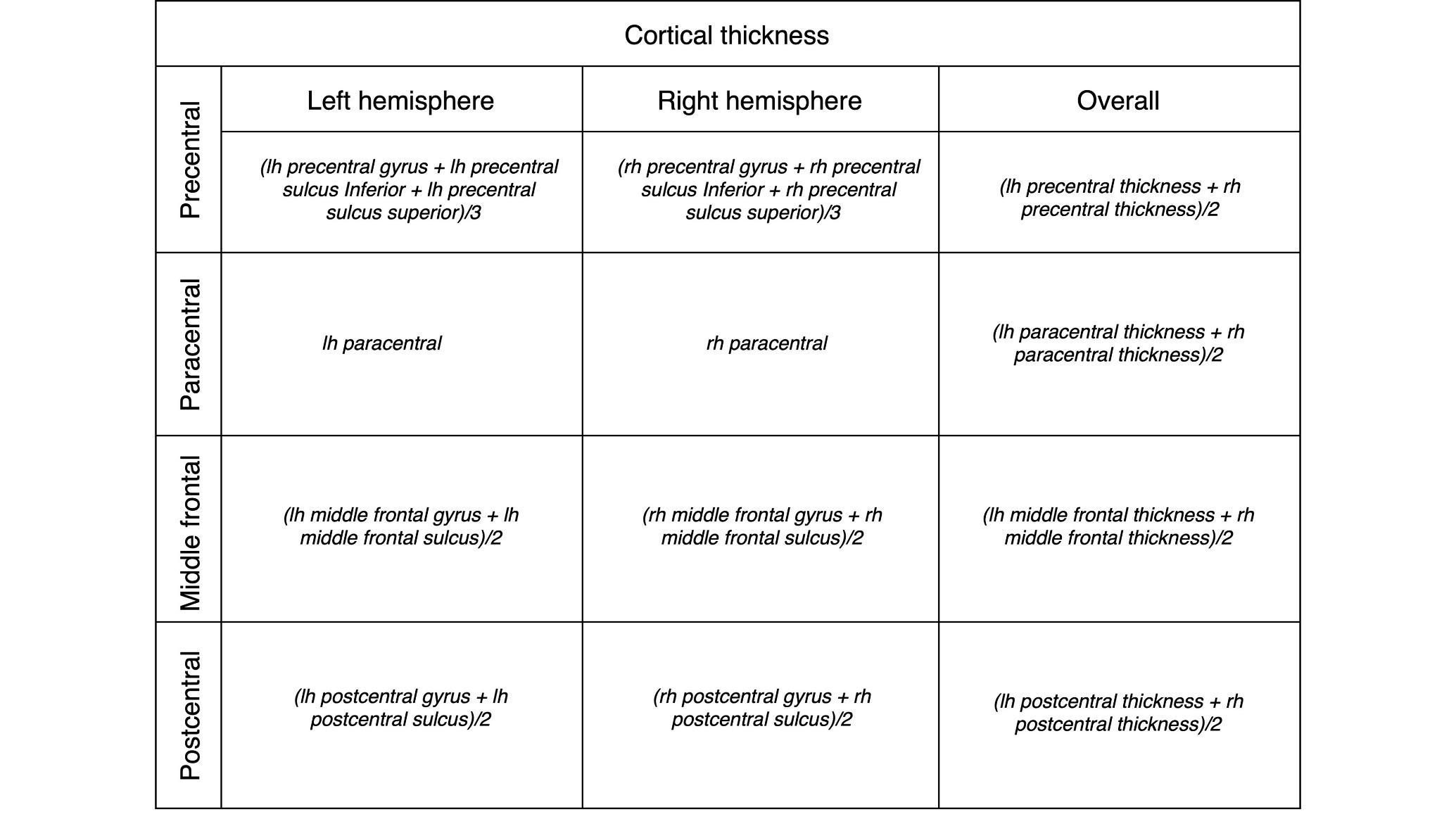

After calculating the thickness and volume data, the distribution for each variable was plotted. We identified and excluded the outliers from further analysis. The final distributions of volume and thickness data are shown in Figures S1 and S3, respectively. The boxplot distribution of volume and thickness data are shown in Figures S2 and S4, respectively.

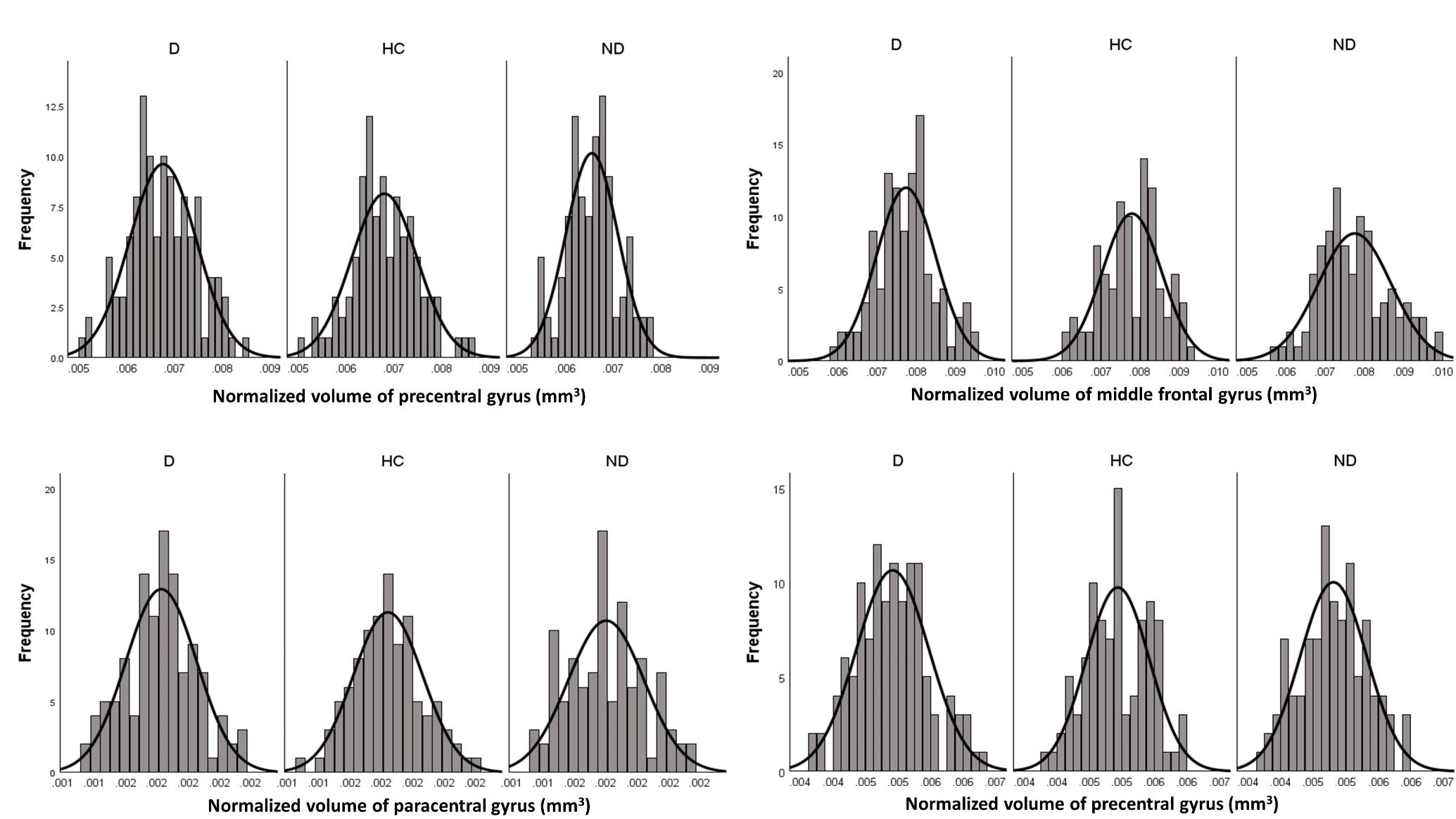

**Figure S1: Histograms representing each group's volumetric distribution of the four cortical ROIs**. D: Dyskinetics, HC: Healthy Controls, ND: Non-Dyskinetics.

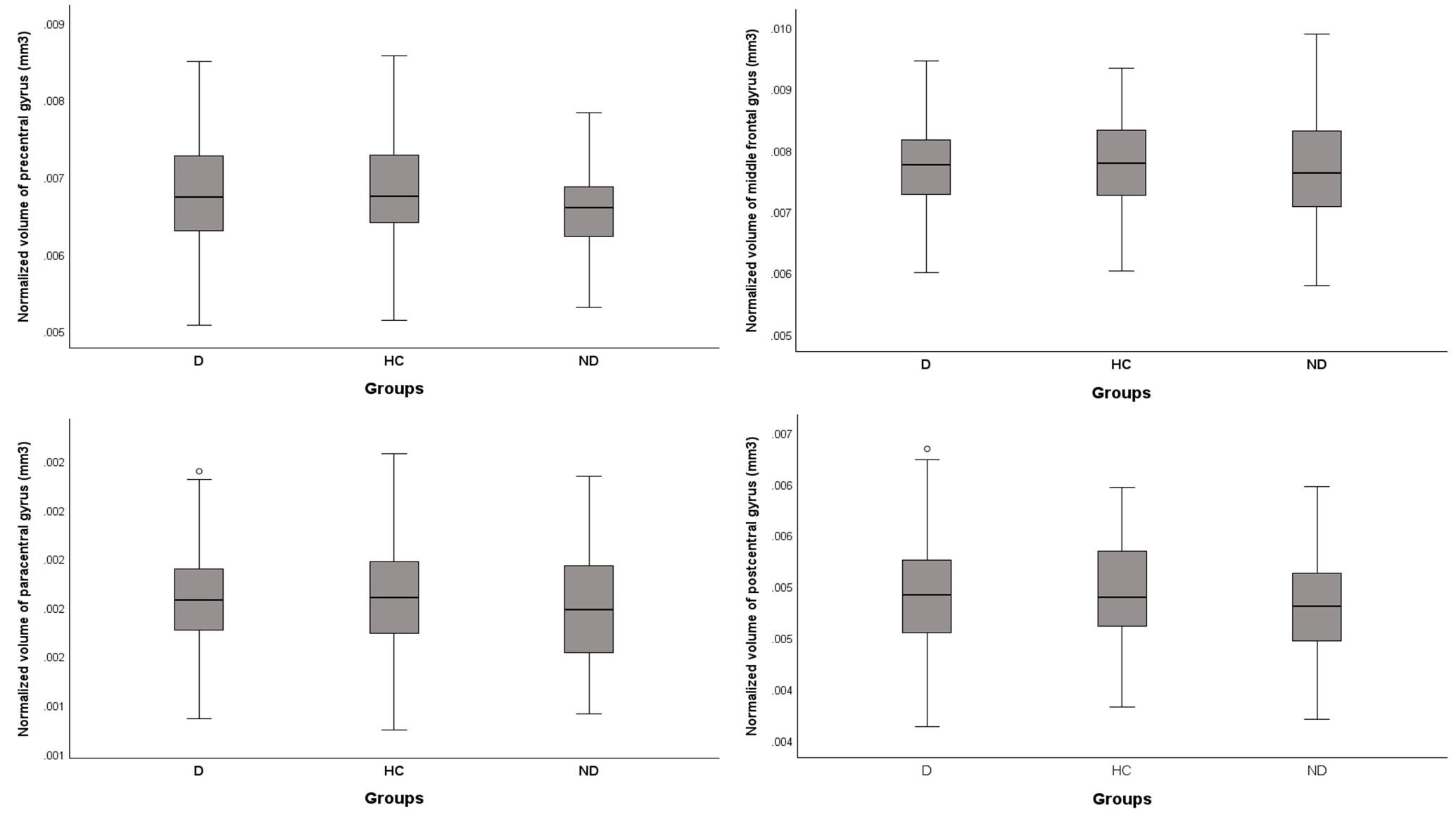

**Figure S2: Boxplots representing each group's volumetric distribution of the four cortical ROIs**. D: Dyskinetics, HC: Healthy Controls, ND: Non-Dyskinetics.

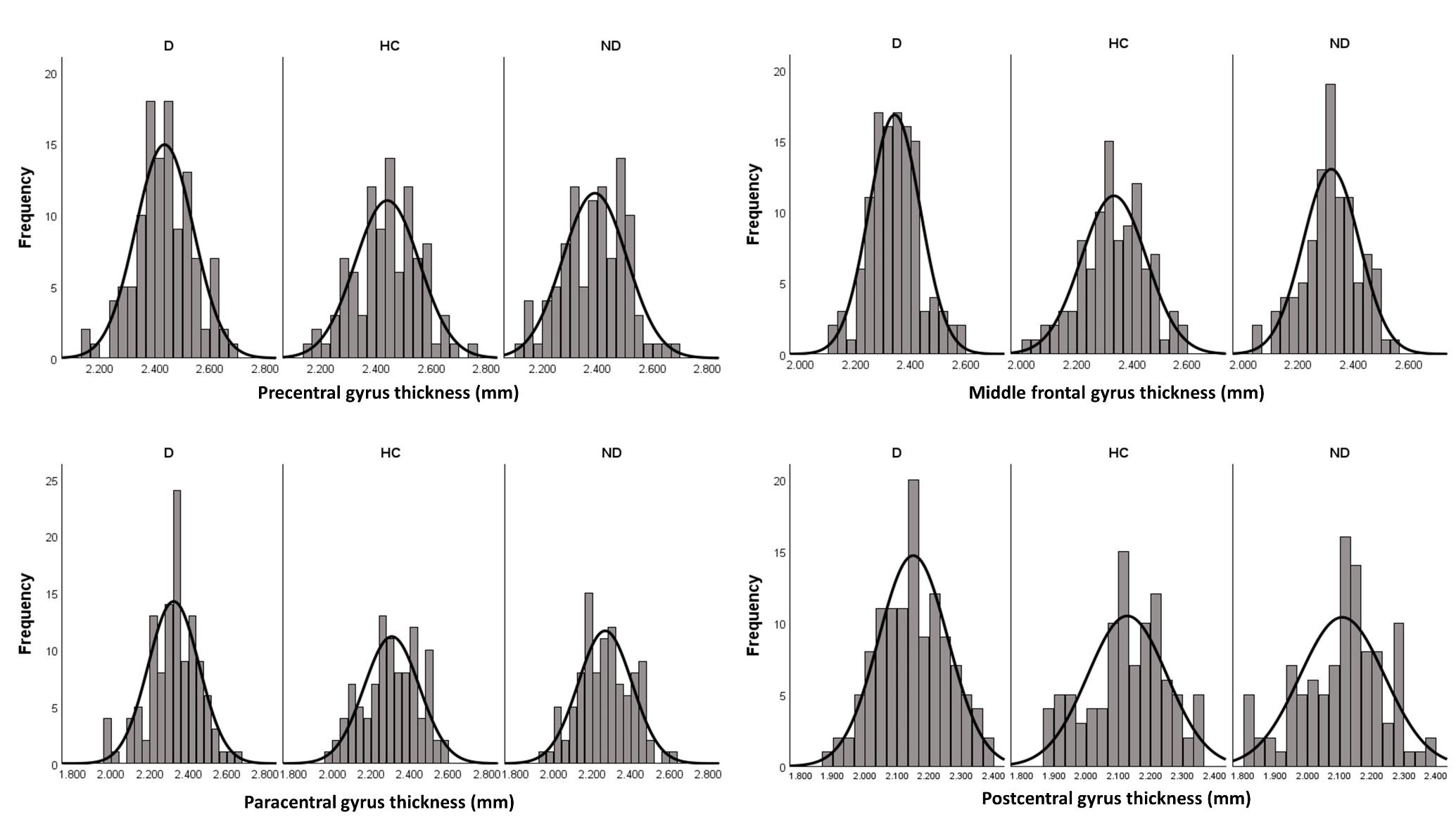

**Figure S3: Histograms representing each group's thickness distribution of the four cortical ROIs.** D: Dyskinetics, HC: Healthy Controls, ND: Non-Dyskinetics.

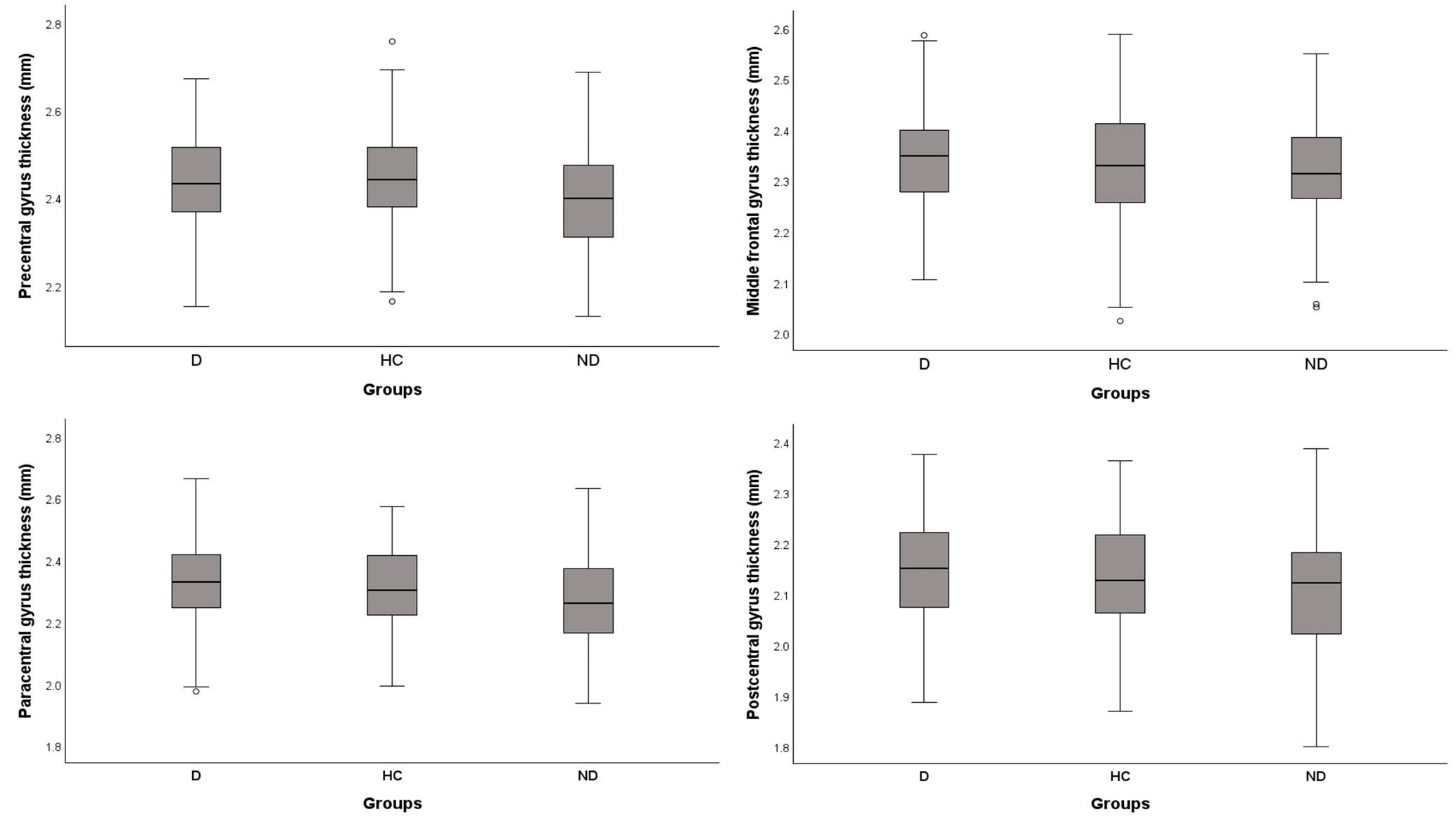

**Figure S4: Boxplots representing each group's thickness distribution of the four cortical ROIs.** D: Dyskinetics, HC: Healthy Controls, ND: Non-Dyskinetics

**Subcortical volumes**

For 5 subcortical ROIs, we averaged the normalized volumes of each region across the hemispheres. We tested the distribution of the data across the groups. We identified and excluded the outliers from further analysis. The final distribution of the subcortical volumes is shown in Figures S5 and S6.

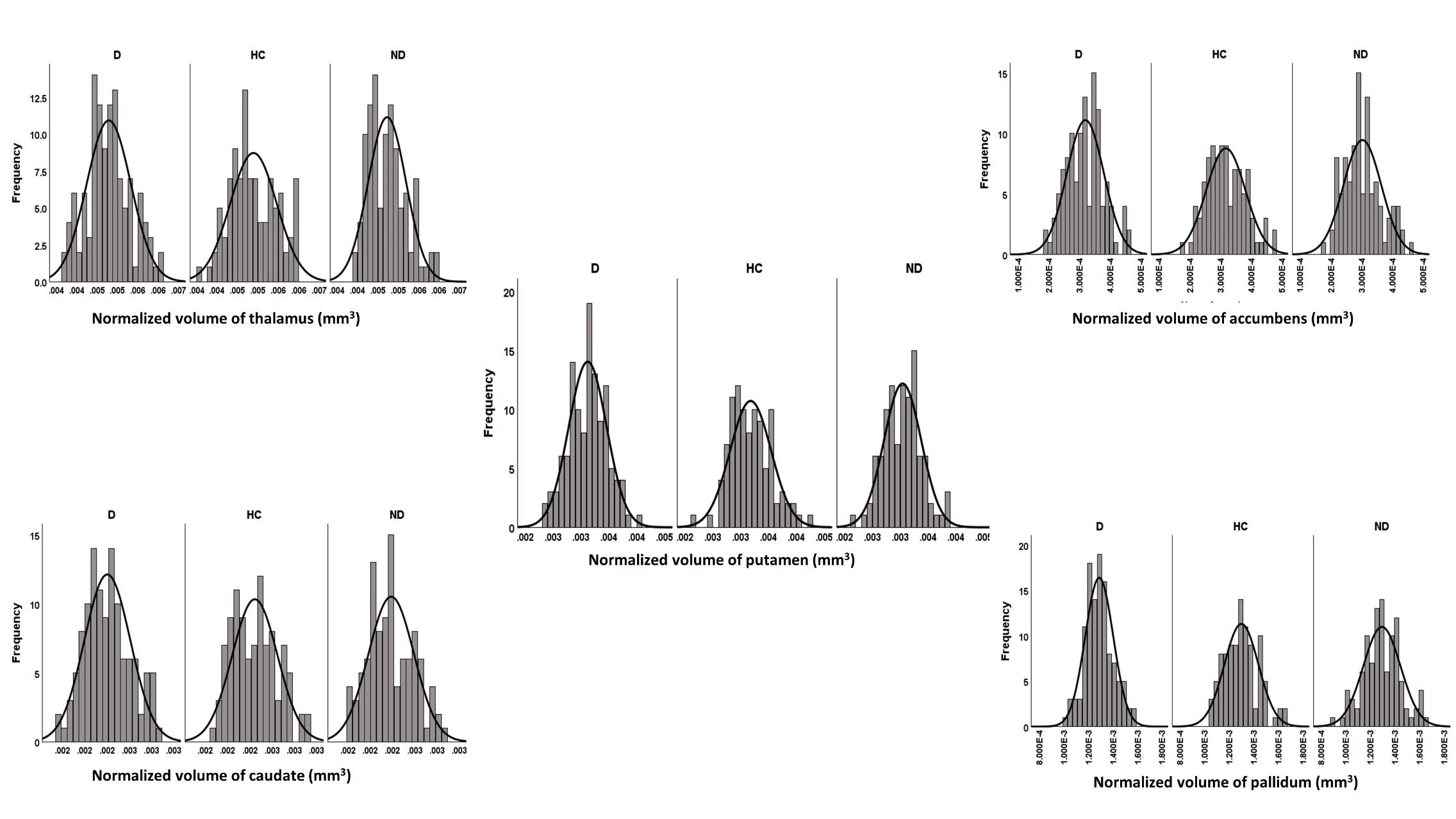

**Figure S5: Histograms representing each group's normalized volumetric distribution of the five subcortical ROIs.** D: Dyskinetics, HC: Healthy Controls, ND: Non-Dyskinetics.

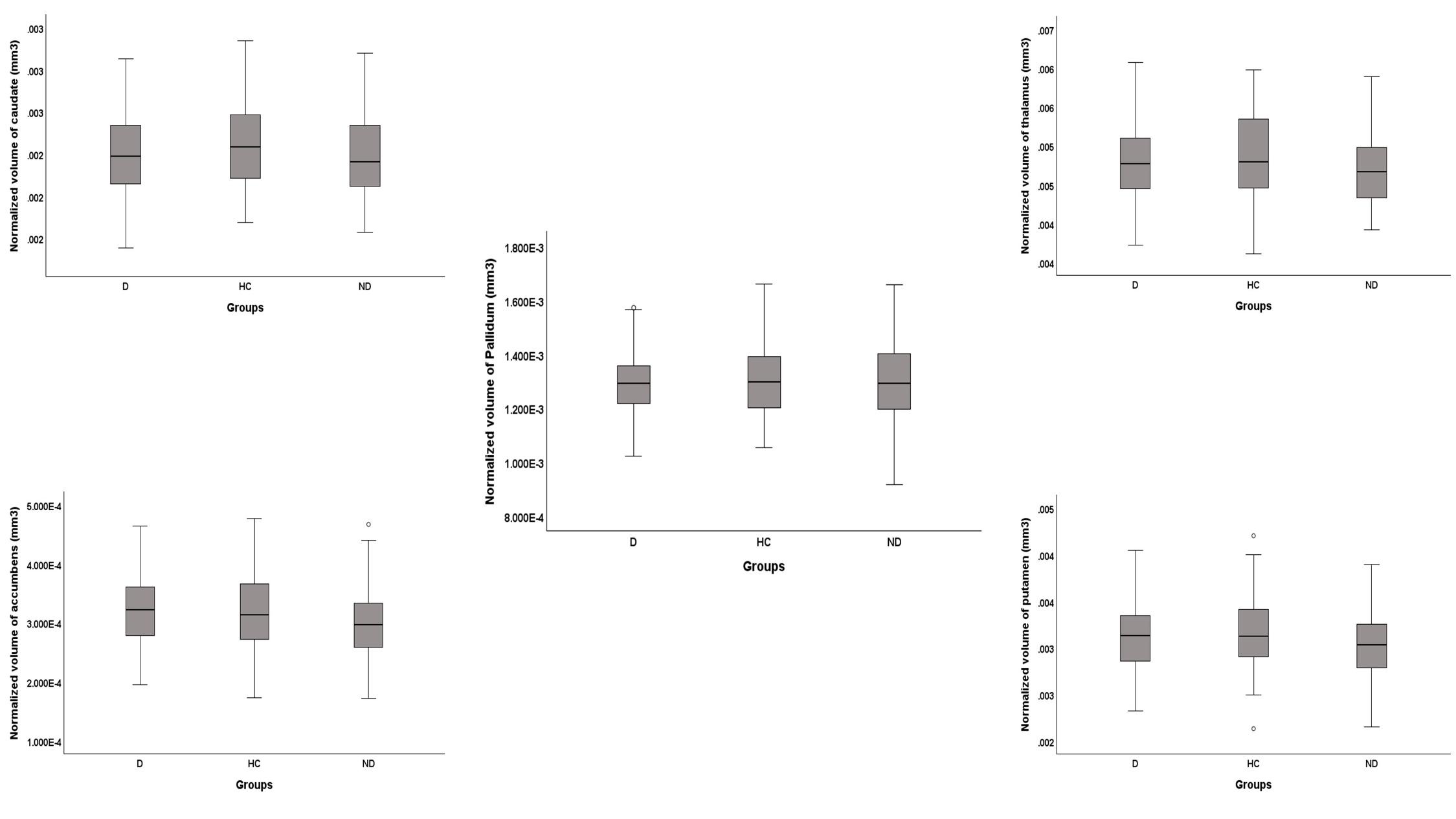

**Figure S6: Boxplots representing each group's normalized volumetric distribution of the five subcortical ROIs.** D: Dyskinetics, HC: Healthy Controls, ND: Non-Dyskinetics.

**Scalar Projection Values**

The distribution of the 10 variables (5 subcortical regions per hemisphere) is shown in Figures S7 and S8.

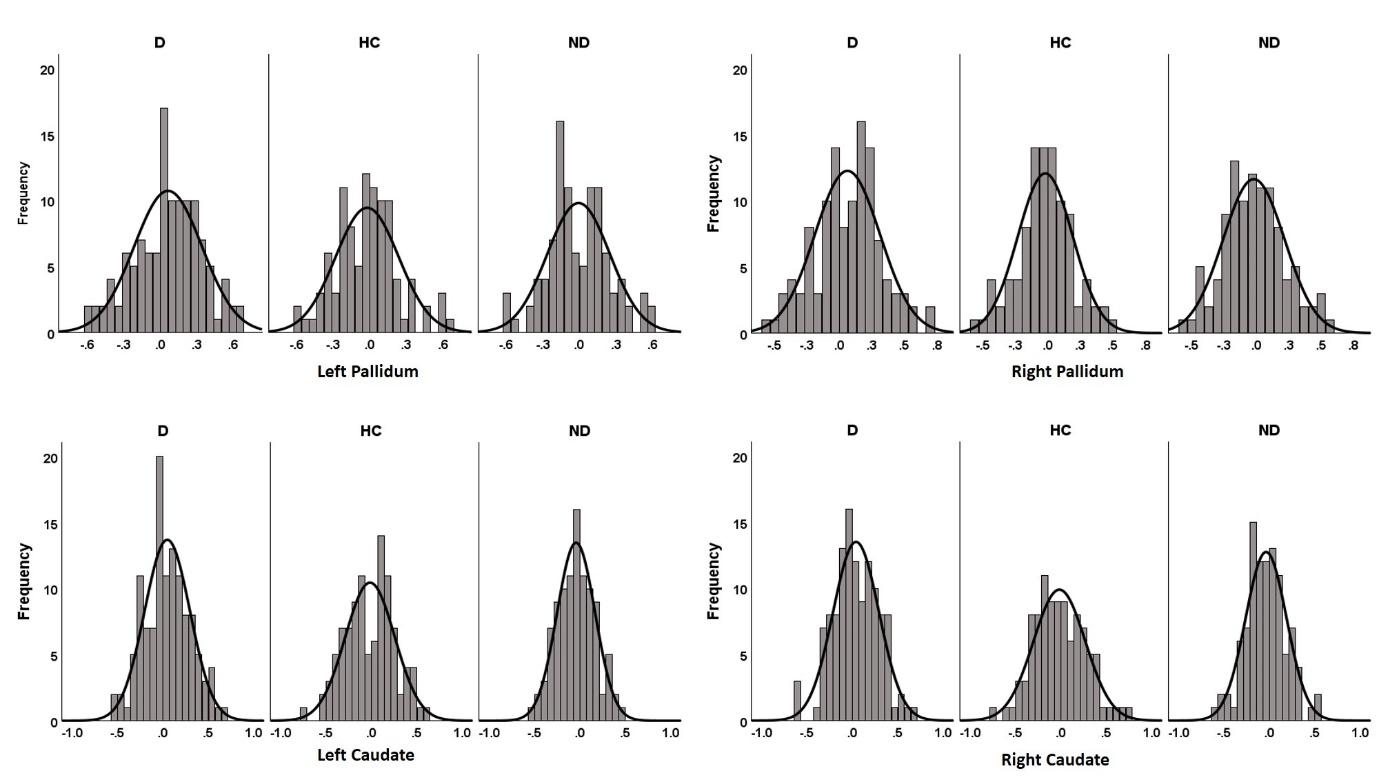

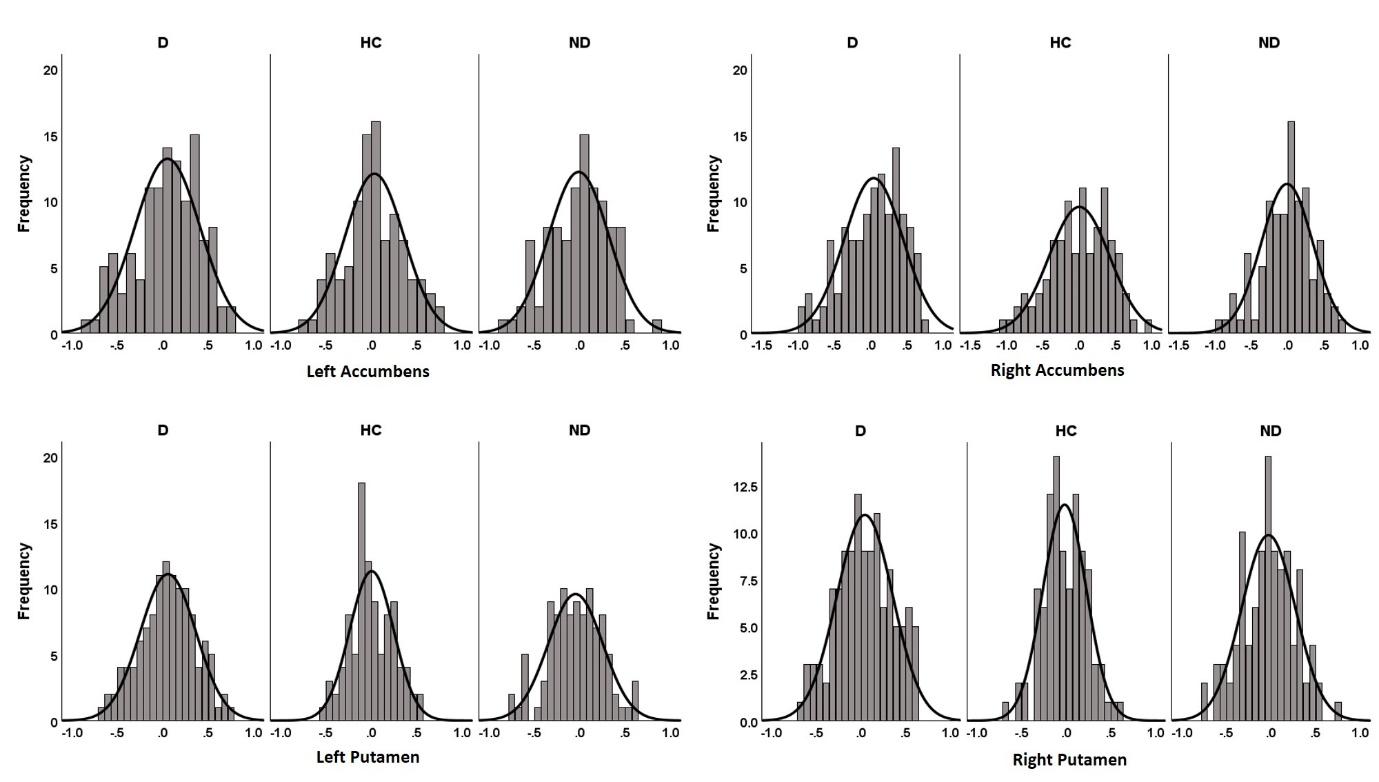

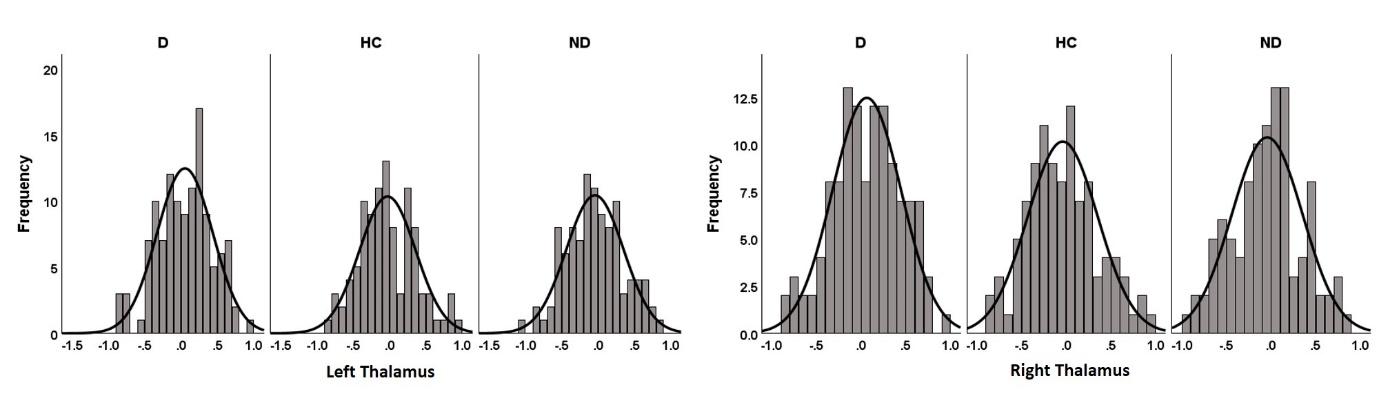

**Figure S7: Histograms representing the distribution of scalar projection values for the 10 subcortical ROIs in each group.** D: Dyskinetics, HC: Healthy Controls, ND: Non-Dyskinetics.

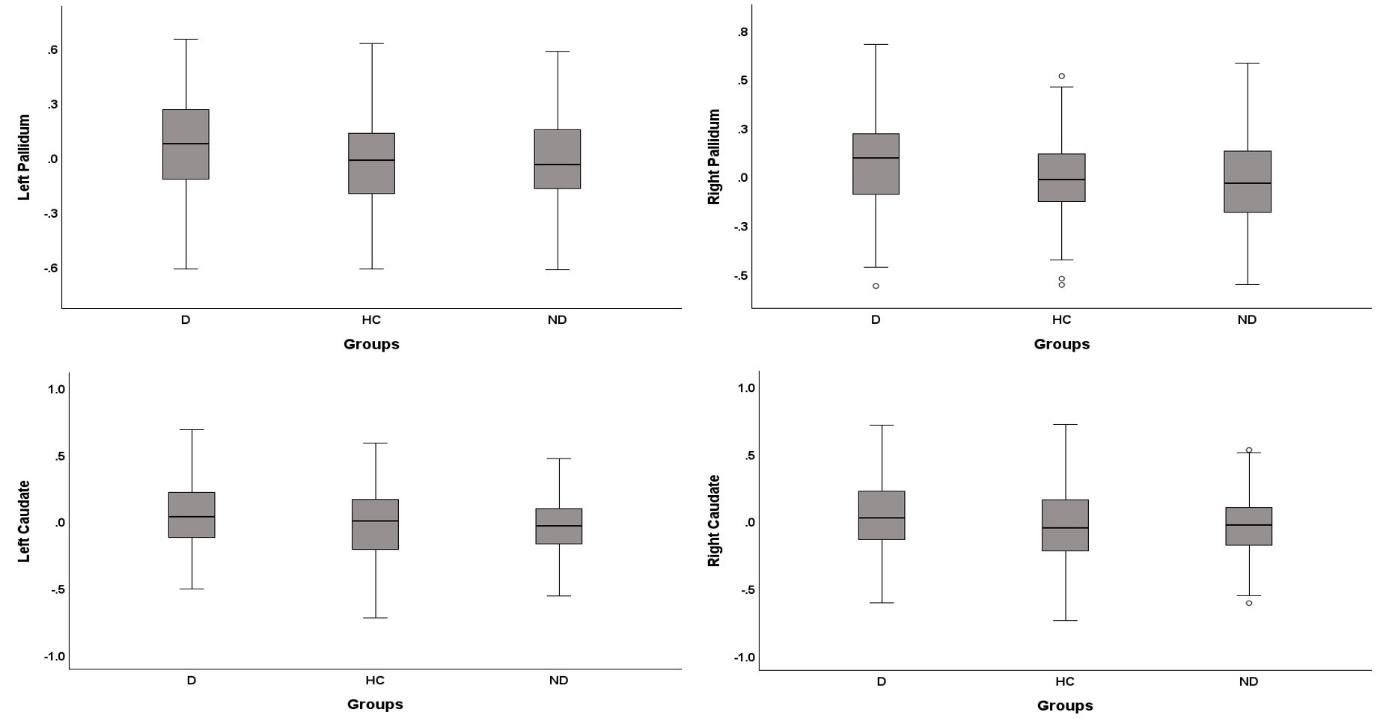

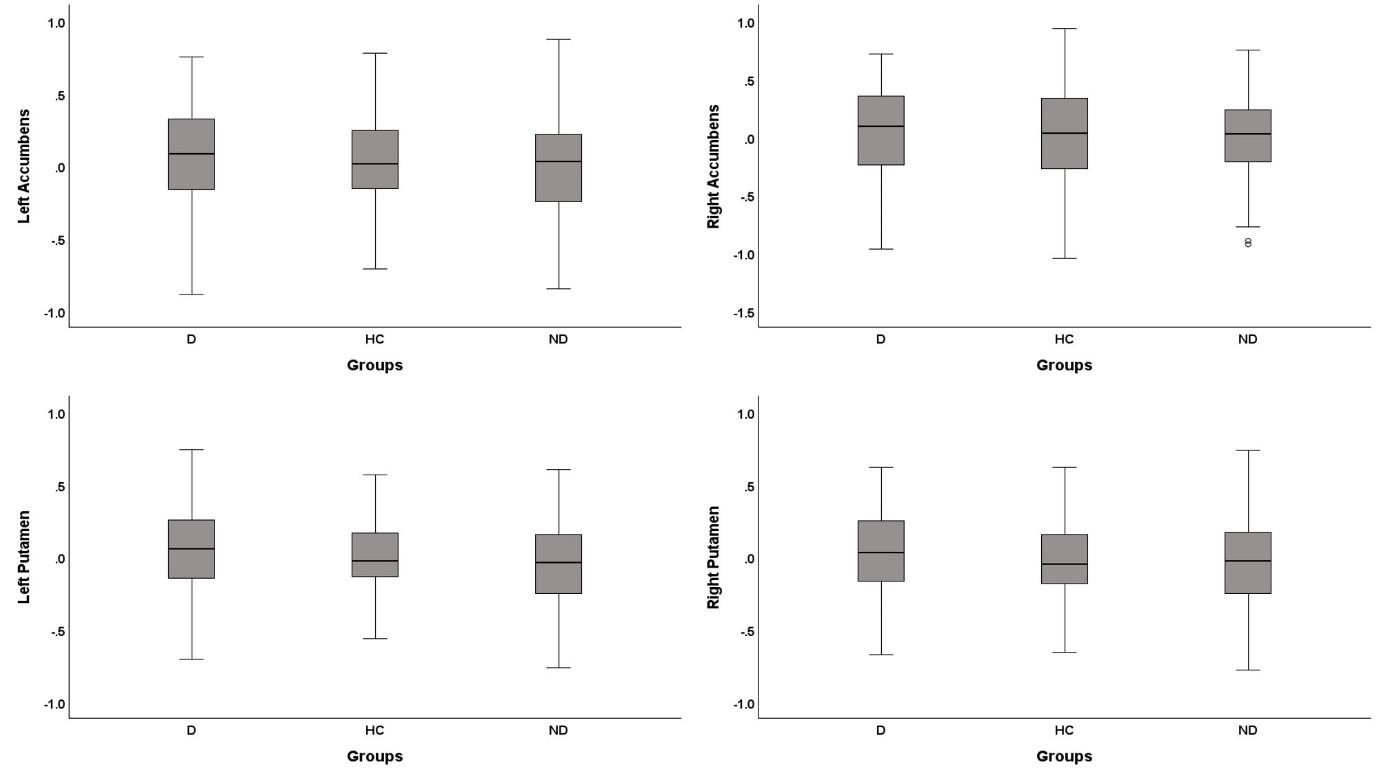

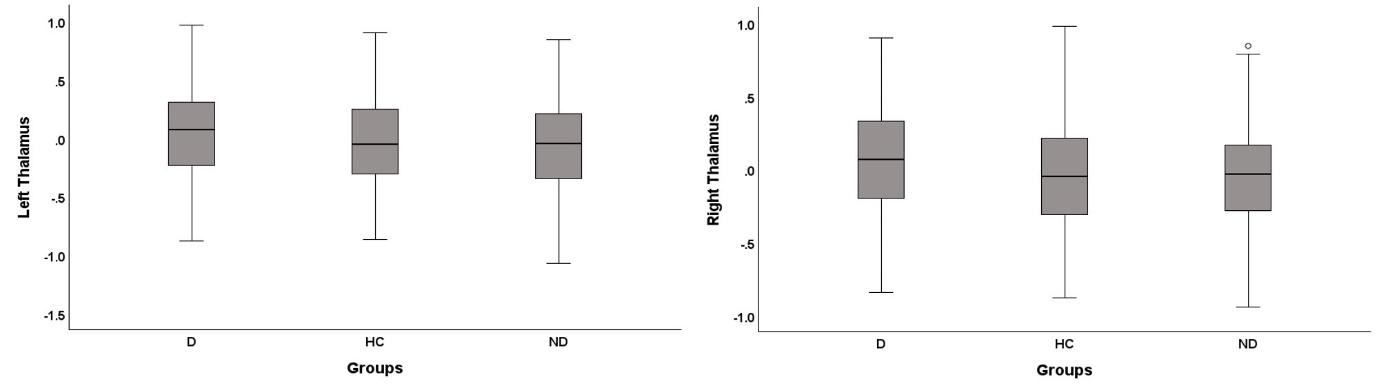

**Figure S8: Boxplots representing the distribution of scalar projection values for the 10 subcortical ROIs in each group.** D: Dyskinetics, HC: Healthy Controls, ND: Non-Dyskinetics.

**Resting-state functional MRI data**

**Medication status**

While most patients started taking medicine by the time of fMRI acquisition, a subset did not. Those under medication were stratified into either the ON or OFF state. However, some patients had an undetermined medication status at the time of scanning, thus classified as 'unknown'. Analysis employing a chi-square test demonstrated no statistically significant variance in medication status between individuals experiencing dyskinetic symptoms and those without (refer to Table S5).

**Table S5: Medication status of D and ND patients during fMRI scanning.** D: Dyskinetic; ND: Non-dyskinetic.

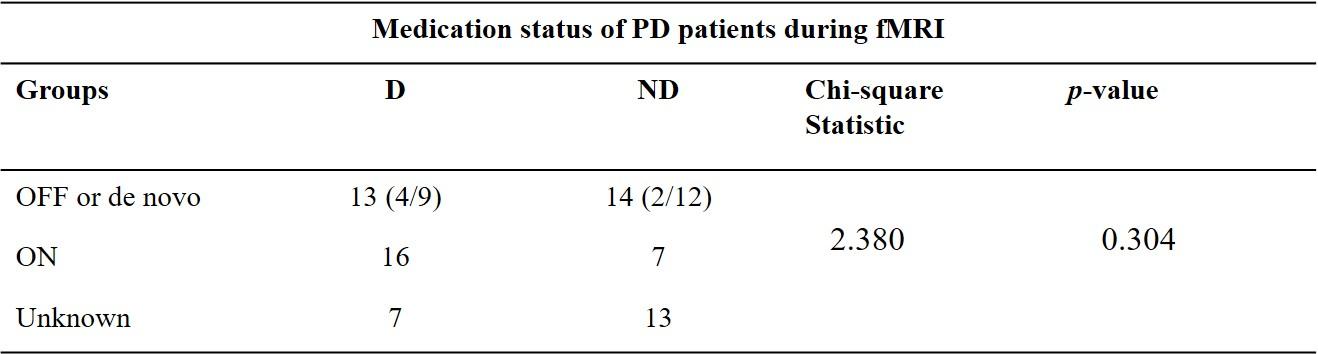

**Quality assurance report**

After preprocessing the rs-fMRI data, we looked into the following quality check measures for all subjects before any further analysis: Number of valid (non-outlier) scans, number of outlier scans, maximum motion, mean motion, BOLD signal standard deviation (after denoising), global correlation at rest. Violin plots in Figure S9 show the distribution of these measures for our dataset. We also checked the distribution of functional connectivity values before and after denoising (Figure S10).

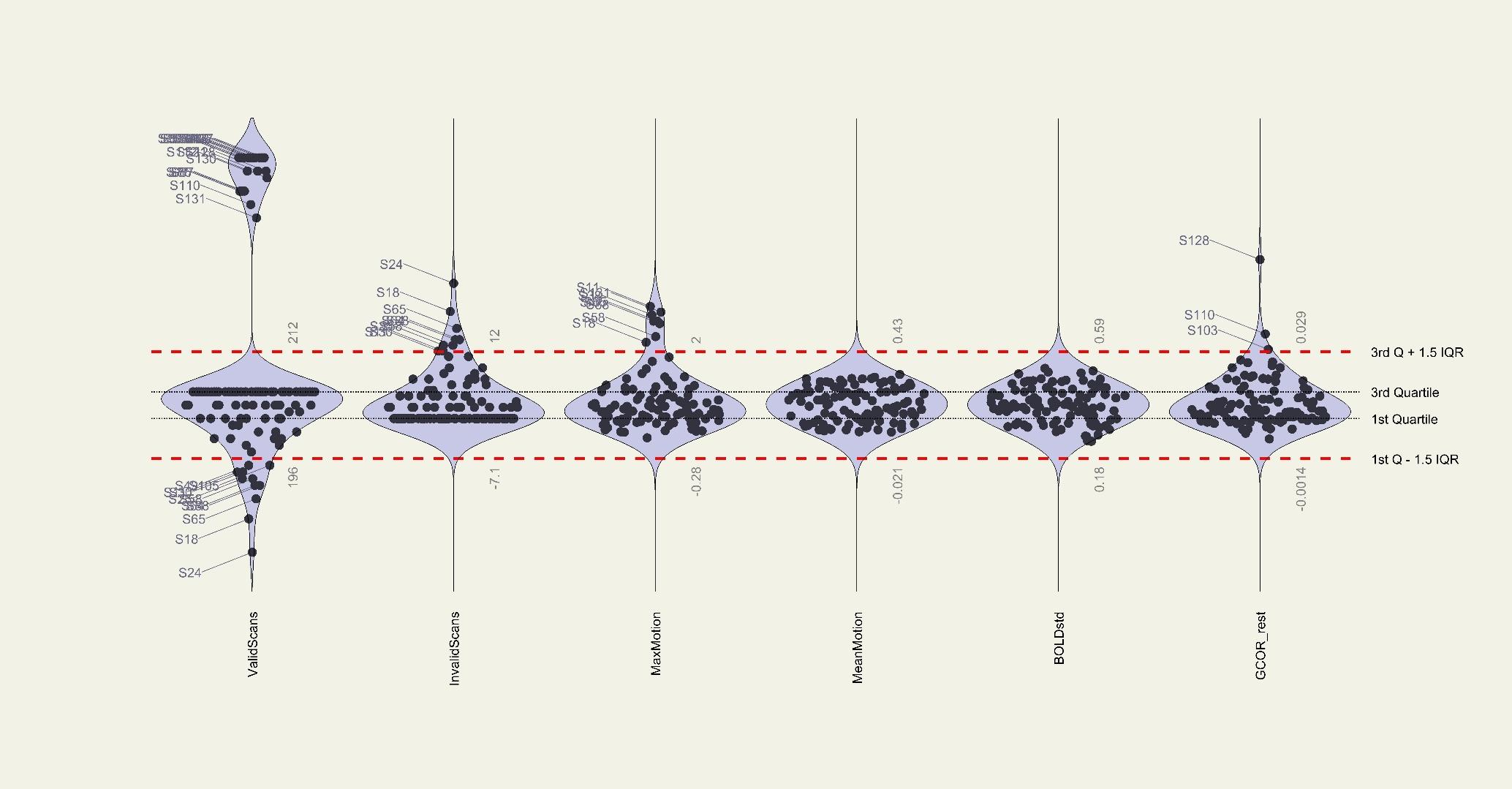

**Figure S9: Violin plots showing the distribution of quality check measures for the dataset.**

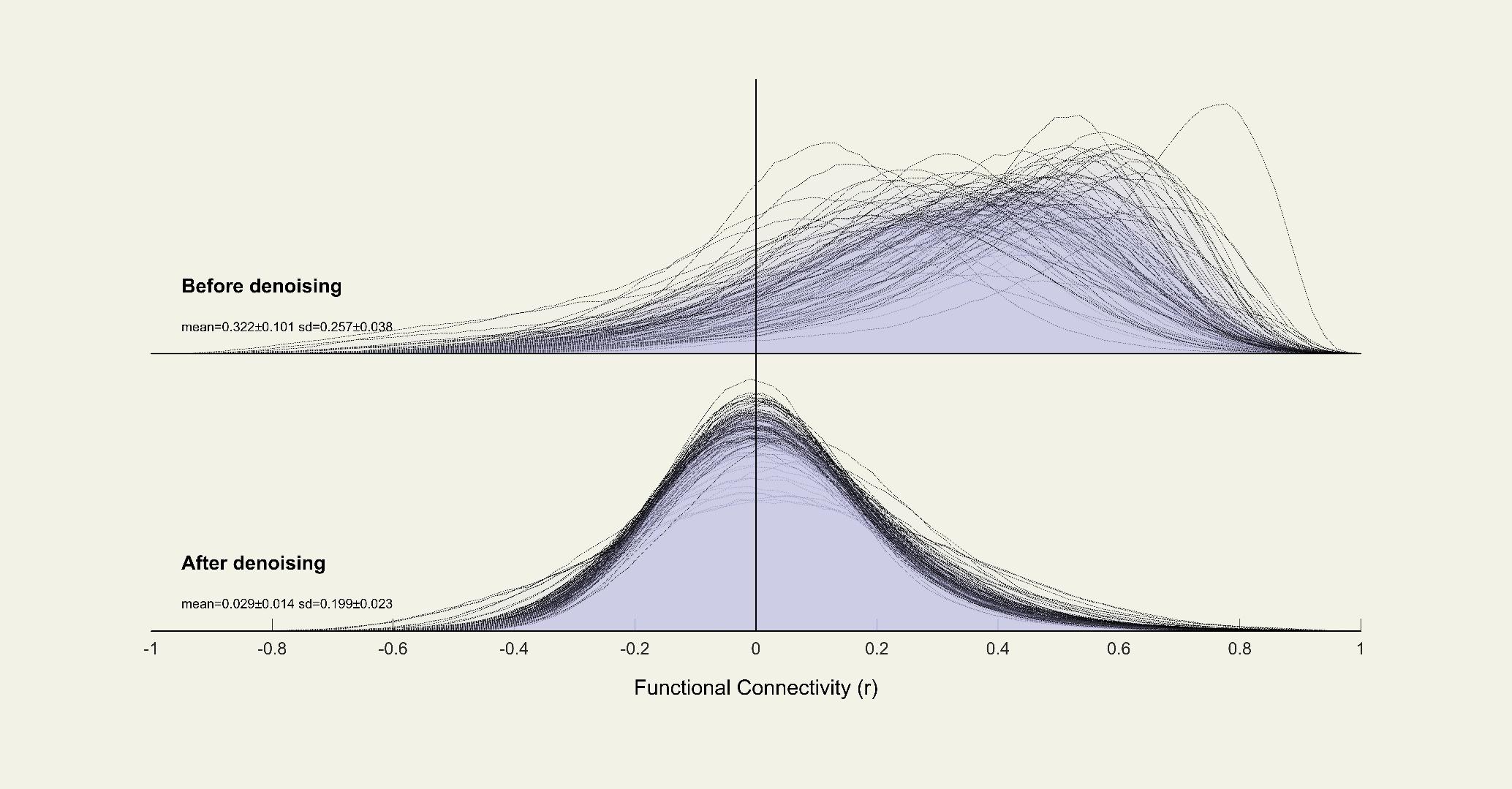

**Figure S10: The distribution of functional connectivity values before and after denoising.**

**Results**

*Whole-Brain Exploratory Analysis*

The exploratory analysis was done by creating the FreeSurfer Group Descriptor (FSGD) files (3 groups) and contrasts. First, we concatenated all subjects’ data into a single dataset and resampled it to fsaverage template using the mri_preproc command. We used a smoothing kernel with FWHM=10mm. Then we fitted a GLM with the mri_glmfit command to look for any group differences. We corrected for multiple comparisons using Monte Carlo simulations with a significance threshold of 1.3, corresponding to p < 0.05.

In the baseline data analysis, we found a significantly higher volume of left posterior cingulate and fusiform gyrus in dyskinetic patients compared to healthy controls (Table S6). However, the exploratory analyses did not reveal any significant differences in the cortical thickness, area, or volume between dyskinetics vs non-dyskinetics or non-dyskinetics vs healthy controls.

**Table S6: Summary table of the exploratory analysis showing significant clusters between D and HC groups** with their respective MNI coordinates, cluster size, and CWP (cluster-wise p values).

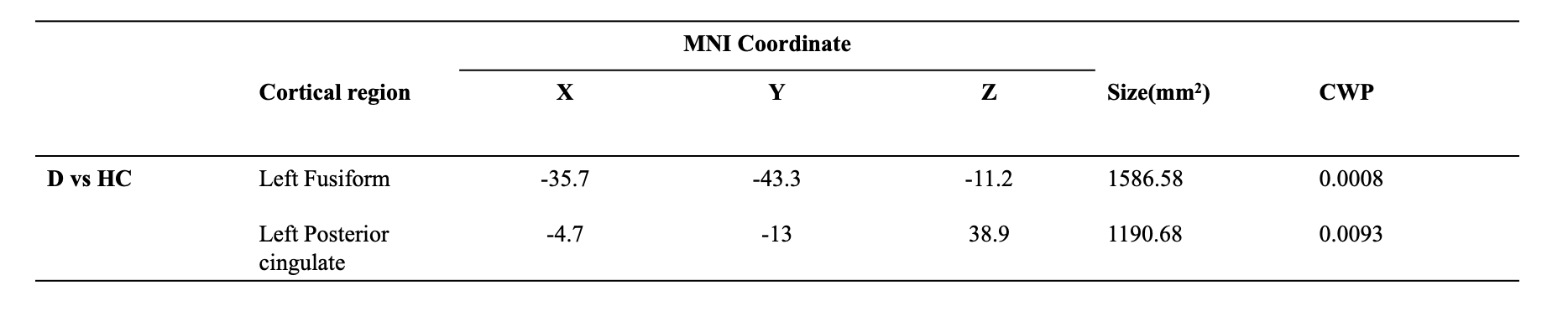

*Vertex-based analysis*

We analysed shape differences between dyskinetics and non-dyskinetics after controlling for ‘Age’ and ‘Dyskinesia-free duration’. We found the shape differences in the left Caudate (Table S7).

**Table S7: Basal ganglia vertex analysis results.**

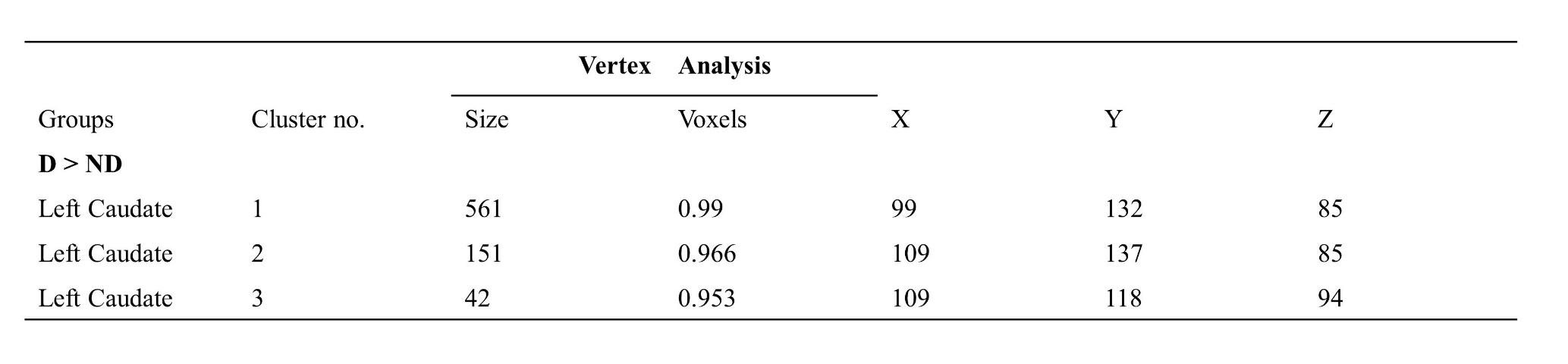
D: Dyskinetics, ND: Non-dyskinetics.

**Effect of Age on subcortical volumes**

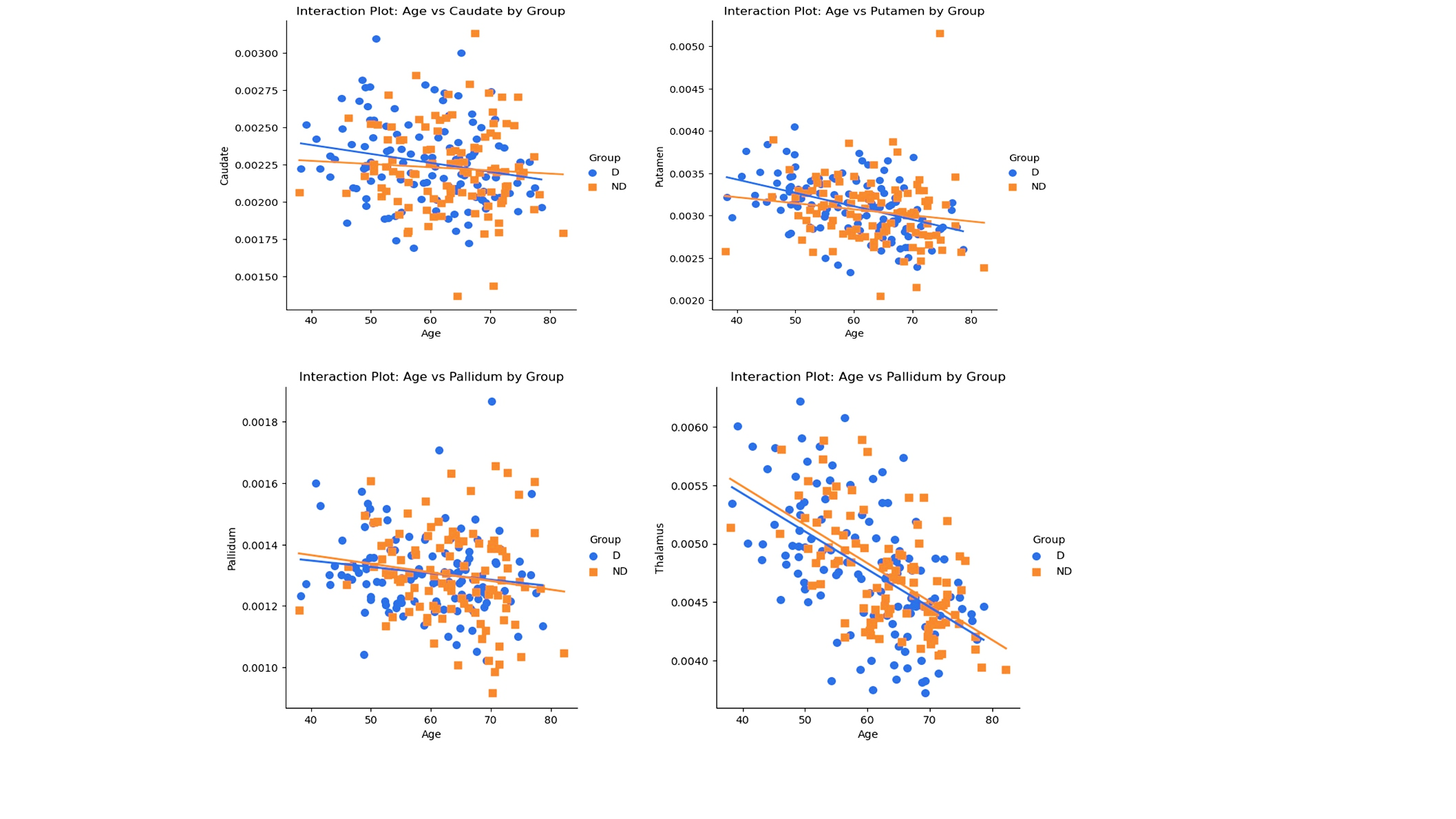

**Figure S13: Scatter plot showing the significant interaction of age with subcortical volumes for dyskinetics (Blue) and non-dyskinetics (Orange).**

**Effect of disease duration**

The disease duration showed a significant effect on cortical thicknesses in precentral, paracentral and middle frontal regions in PD subgroups (Figure S14).

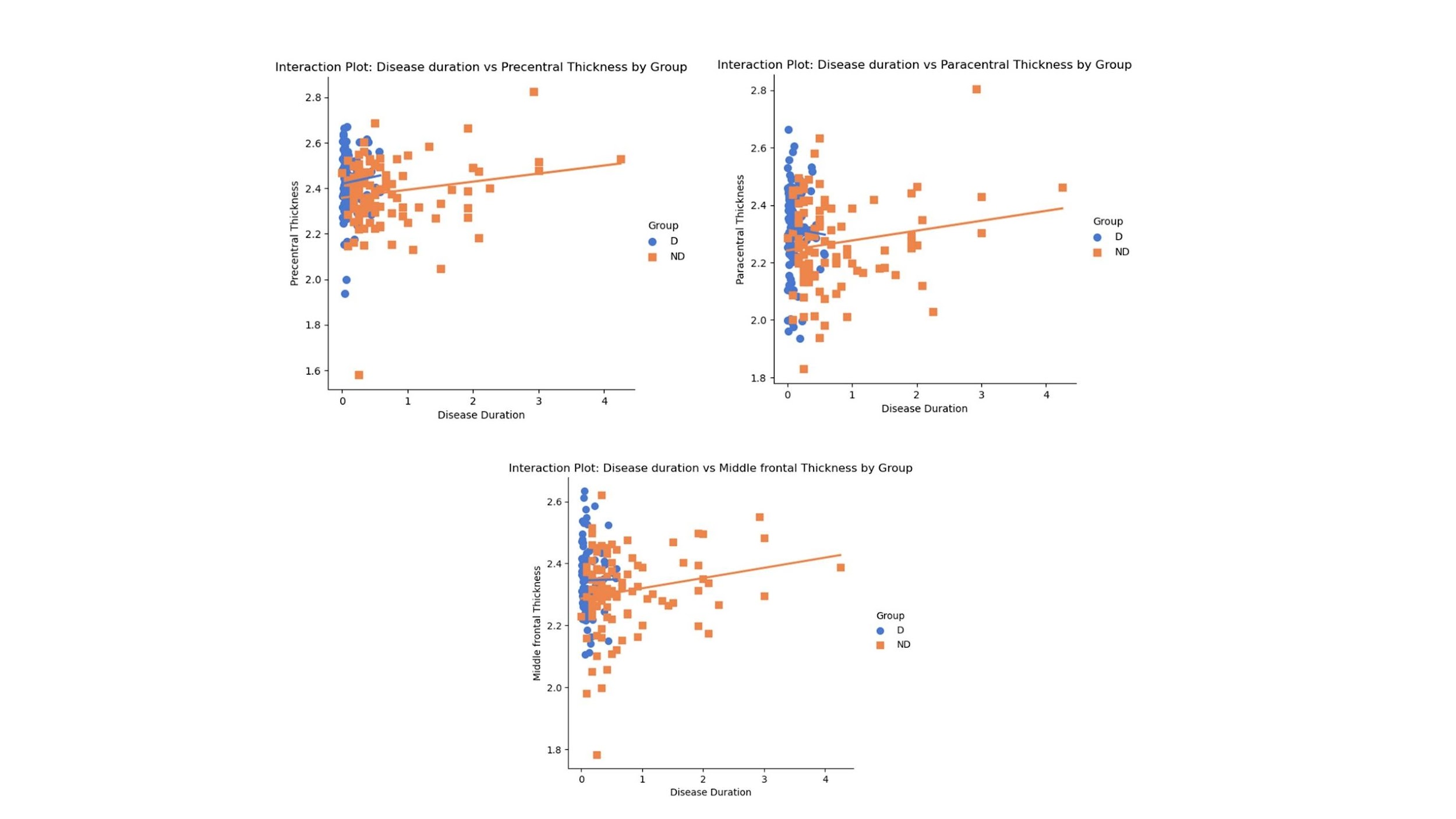

**Figure S14: Scatter plots showing the significant relationship of disease duration with cortical thickness.**

**ROI-to-ROI connectivity analysis**

Additionally, we compared the two subgroups of PD patients after controlling for ‘age’ and ‘dyskineisa-free period’. Our results show the same connectivity differences as shown when we only controlled for ‘age’ (Figure S15).

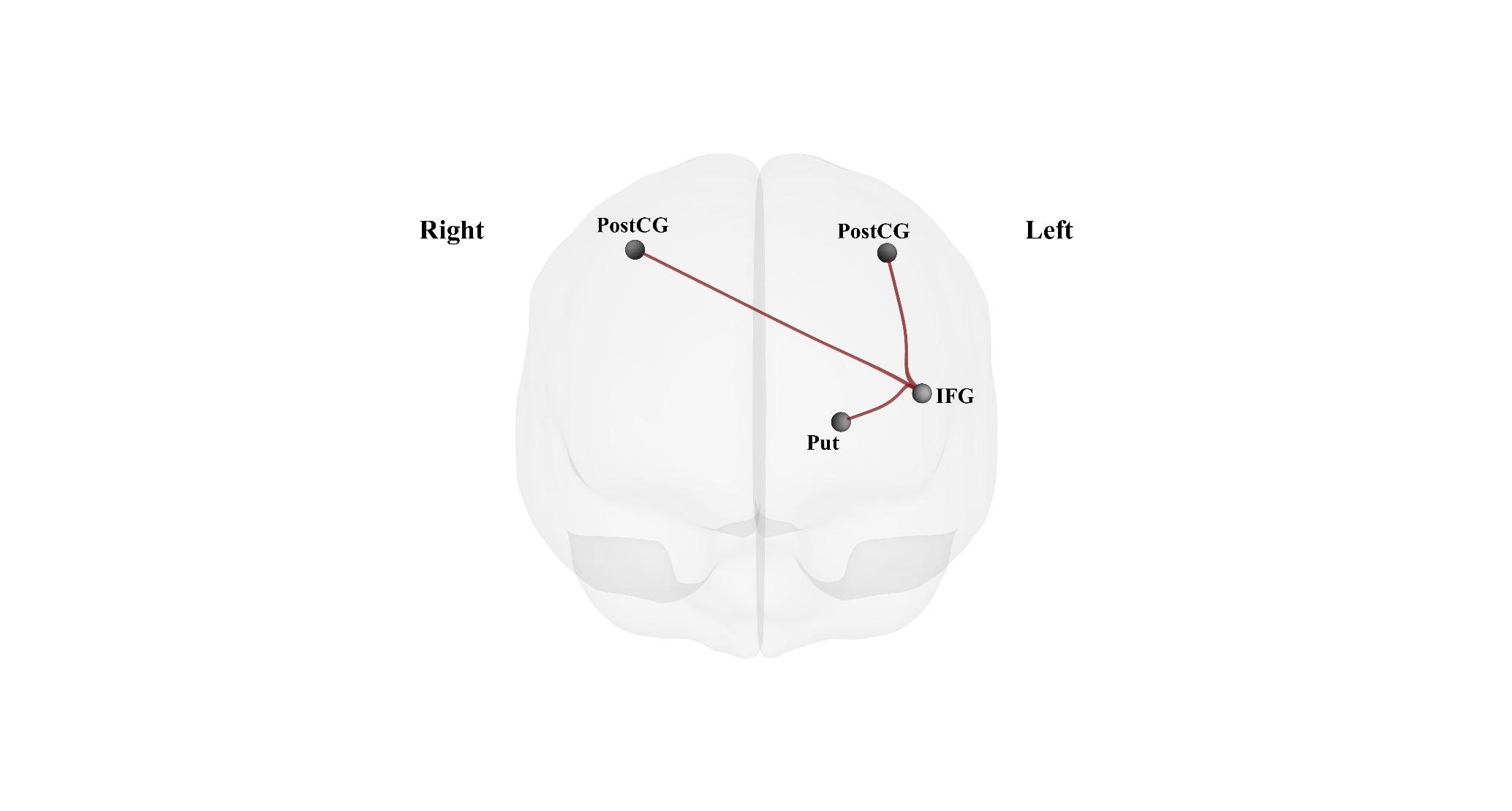

**Figure S15: ROI-to-ROI connectivity differences between dyskinetics and non-dyskinetics with age and dyskinesia-free period as covariates.**
